# MicroRNA-205 promotes hair regeneration by modulating mechanical properties of hair follicle stem cells

**DOI:** 10.1101/2022.12.29.522250

**Authors:** Jingjing Wang, Yuheng Fu, Wenmao Huang, Ritusree Biswas, Avinanda Banerjee, Joshua A. Broussard, Zhihai Zhao, Dongmei Wang, Glen Bjerke, Srikala Raghavan, Jie Yan, Kathleen J. Green, Rui Yi

## Abstract

Stiffness and actomyosin contractility are intrinsic mechanical properties of animal cells required for the shaping of tissues. However, whether tissue stem cells (SCs) and progenitors located within SC niche have different mechanical properties that govern their size and functions remains unclear. We show that hair follicle SCs in the bulge are stiff with high actomyosin contractility and resistant to size change, whereas hair germ (HG) progenitors are soft and periodically enlarge and contract during quiescence. During activation, HGs reduce contraction and more frequently enlarge, a process that is associated with weakening of the actomyosin network, nuclear YAP accumulation and cell cycle re-entry. Induction of miR-205, a novel regulator of the actomyosin cytoskeleton, reduces actin contractility, decreases the stiffness of the bulge and HG, and activates hair regeneration in young and old mice. This study reveals the control of tissue SC size and activities by spatiotemporally compartmentalized mechanical properties and demonstrates the possibility to stimulate tissue regeneration by fine-tuning cell mechanics.

## Introduction

Tissue regeneration is essential for homeostasis, wound repair and aging. Adult stem cells (SCs) and their microenvironment control tissue regeneration by constantly sensing and responding to extrinsic and intrinsic cues ranging from signaling molecules to mechanical forces. The response of living cells to mechanical forces is modulated by physical properties of both external microenvironment, such as extracellular matrix (ECM) stiffness, and internal force-generating machinery, such as the cytoskeleton (1, 2). These mechanical properties have a profound impact on cell behaviors including cell adhesion, division, migration and differentiation (1, 3, 4). However, although mechanical forces continuously regulate cellular and tissue functions, it has been challenging to directly observe their effects on cell size and activities of SCs within the SC niche during homeostasis and regeneration. It is unclear whether spatiotemporally demarcated SC activities, such as quiescence and cell cycle re-entry, within distinct regions of the SC niche is controlled by local stiffness and differentially generated mechanical forces.

Mammalian skin is an excellent experimental system to interrogate the function of mechanical mechanisms on SC activity, tissue regeneration, wound repair and tumorigenesis (5–10). A subset of epidermal SCs rapidly responds to external force-induced stretch by transiently activating cell division (5). Perturbations in the force-generating cytoskeleton and ECM through genetic deletion of key components of these machinery change mechanical properties of the cells (6, 8), and these changes compromise tissue functions. Interestingly, basal epithelial SCs and cultured keratinocytes have been shown to control their cell-cycle progression in a cell size-dependent manner (11, 12). However, whether cell mechanics controls cell size and regulates the transition between quiescence and activation of hair follicle stem cells (HF-SCs) and HG progenitors remains poorly understood. In this study, we uncovered spatiotemporally demarcated mechanical properties of tissue SCs and progenitors as tunable features to promote hair regeneration through the control of cell size dynamics.

## Results

### HF-SC compartment is demarcated by differential mechanical stiffness and actomyosin contractility

To determine cell dynamics within the HF-SC compartment, including both the bulge and HG (13, 14), during quiescence and activation, we performed non-invasive intravital imaging (15, 16). We monitored HF cells labelled with H2b-GFP (*Krt14-H2b-GFP* mice) beginning from late catagen (~postnatal day 19, P19) to early anagen (~P27). The size and position of individual HF-SCs in the bulge remained largely constant over the period. However, the size and shape of HG, usually consisting of ~10-20 stem and progenitor cells, showed substantial (~35%) changes without cell proliferation or death during the same period (Fig. 1A and fig. S1, A and B). To ensure that H2b-GFP nuclear signals accurately measured the HG volume, we used *Krt14-H2b-GFP/Rosa-mTmG* to mark the nuclei and cell membrane, respectively. Double labeling revealed that stem and progenitor cells in the bulge and HG of HFs comprised mainly the nuclei with minimal cytoplasm such that the HG size quantified by H2b-GFP or membrane-localized tdTomato was similar for telogen and early anagen HGs (fig. S1, C and D). Because H2b-GFP distinguished individual cells and cell division events, and very low laser power was required for imaging H2b-GFP, we used H2b-GFP in subsequent studies to minimize phototoxicity. As shown in Figure 1A, the HG contained the same number of cells but changed the volume, in contrast to the largely invariable bulge, from P19 (late catagen) to P24 (late telogen). For each HG, we designated the largest volume during observation as 100%. The smallest size reached ~54.56% of the largest volume (fig. S1B), usually during the middle of telogen. When HFs transitioned toward anagen regeneration and cell cycle re-entry, the HG size increased (Fig. 1A and fig. S1B). The average size of HG cells ranged from 454 μm^3^ to 735 μm^3^ (fig. S1B).

**Figure 1.**
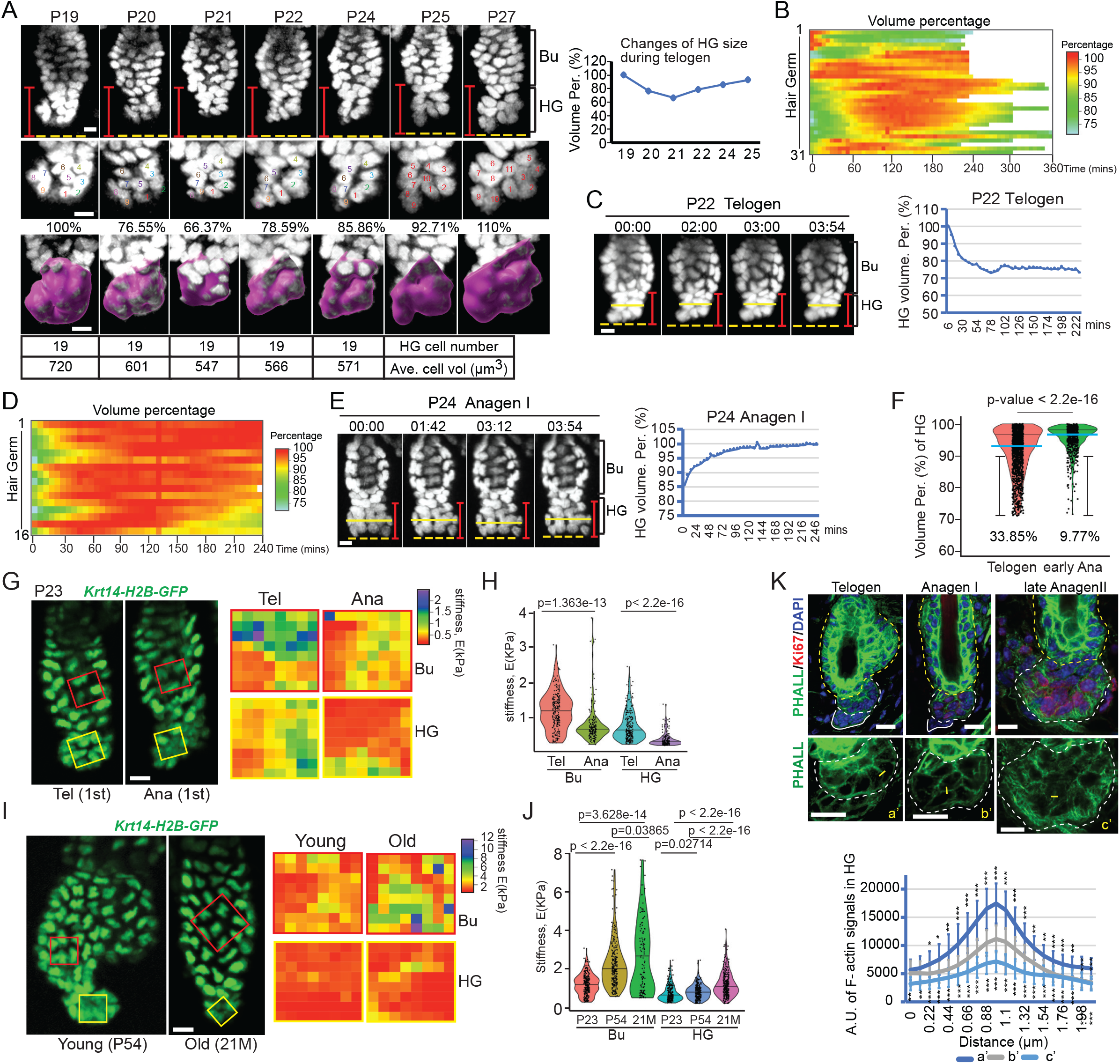
HF-SC compartment is mechanically active and exhibits differential mechanical stiffness and actomyosin contractility during activation. **(A)** Hair germ (HG) showed dynamic volume change in late catagen and telogen in the absence of cell division. Bu, bulge; HG, hair germ. Purple surfaces indicate HG areas used for quantification in Imaris. 65 hair follicles from 4 animals were tracked. (**B)** HGs in telogen showed pulsatile contraction and enlargement. 31 hair follicles from 9 animals were used to calculate volume changes over 4-6 hours. (**C)** An example of HG undergoing continuous contraction (left) with quantification (right) over 4 hours in telogen. (**D)** HGs in early anagen with reduced contraction, compared to HGs in telogen. 16 hair follicles from 5 animals were recorded for quantification. (**E)** An example of HG undergoing continuous enlargement (left) with quantification (right) over 4 hours in early anagen without cell division. (**F)** HGs in early anagen showed reduced contraction, compared with those in telogen. The light blue lines represented median (med), and black lines in violin plot represented the third quarter of all time points. **(G)** Stiffness of bulge and HG in first Telogen and Anagen was measured ex vivo by atomic force microscopy (AFM). Force maps (with 64 measured points) of bulge and HG were showed in the right panel. **(H)** Quantification of stiffness (Young’s modulus) of bulge and HG in first Telogen and Anagen. 4 hair follicles from 2 animal in Telogen and Anagen, respectively, were measured. For each area, 64 points were measured. In Telogen, Bu mean= 1196 ± 534 Pa, HG mean= 747.3 ± 414.3 Pa; in Anagen, Bu mean= 798.3 ± 530.7 Pa, HG mean=374.9 ± 183 Pa. **(I)** Stiffness of bulge and HG in young and old back skin was measured ex vivo by AFM. **(J)** Quantification of stiffness (Young’s modulus) of bulge and HG in first Telogen and Anagen, young (P54) and old back skin (26 months old). 5 hair follicles from 2 young animals and old animals were measured. For each area, 64 points were measured. For young hair follicles, Bu mean= 2240.1 ± 1251.1 Pa, HG mean= 824.2 ± 368.0 Pa. For old hair follicles, Bu mean= 2620.9 ± 1868.0 Pa, HG mean= 1207.2 ± 625.2 Pa. (**K)** Cortical F-actin signals were reduced, and cortical cytoskeletons were rearranged during the activation of hair growth. For quantification, mean and s.d. were shown from over 10 line-quantification values (right panel). Scale bar 10 μm in **A, C, E, G, I, K**. P values were determined by Student’s t-test.

To determine the time scale of cell size change observed in HF-SCs and HGs, we next performed intravital time-lapse imaging to quantify the relative size of telogen HFs. We recorded 31 HFs from 9 animals every 6 minutes over 4-6 hours for a total of 1,560 time points during telogen. The bulge HF-SCs remained largely unchanged whereas the HGs were dynamic. For each HG, we designated the largest volume during observation as 100% and calculated the relative volume in each time point such that we could compare the relative HG volumes from different HFs. Three types of HG activities were recorded - 2 out of 31 (6.5%) enlarged continuously, 25 out of 31 (80.6%) HGs dynamically enlarged and contracted, and 4 out of 31 (12.9%) contracted continuously (Fig. 1B and fig. S2A). As an example, one HG continuously contracted and reduced the volume by ~25% over 4 hours during the first telogen at P22 (Fig. 1C and movie S1). Another HG enlarged and contracted intermittently in a 6-hour span (movie S2). The substantial contraction of HG during telogen was further confirmed when we used an *E-cad-CFP* knockin mouse, in which CFP is fused to endogenously expressed E-cad, to visualize adherens junction (AJ) on the cell membrane (fig. S2B and movie S3).

We next examined the HGs during the telogen-to-anagen transition when the new round of hair growth and cell cycle re-entry were initiated. In early anagen, cell division was expected to increase the number of HG cells to fuel hair growth. Indeed, we observed cell division events and used them to mark the beginning of anagen (fig. S2, C and D). Infrequent cell divisions were observed in early anagen, such that there was no more than one cell division in a single HG over the time span of 4-6 hours. Interestingly, these cell divisions did not lead to appreciable increases in HG size (fig. S2D), consistent with experimental observations that cell division is resulted from a larger cell gives rise to two smaller cells (12, 17). Overall, we observed less HG contraction during early anagen, in comparison to telogen HGs. Furthermore, we observed more cases of HG enlargement during the telogen to anagen transition, compared to the telogen phase. In 16 HFs from 5 mice, 4 (25%) HGs exhibited continuous enlargement (movie S4) and only 3 (18.8%) HGs showed substantial (>10%) contraction (Fig. 1, D and E and fig. S2E). Quantification of the relative size of all HGs across 2,215 imaging time points (1,560 time points in telogen and 655 time points in anagen) revealed significantly reduced HG contraction in early anagen (9.77% of all HGs were smaller than 90% of the largest volume) than in telogen (33.85% of all HGs were smaller than 90% of the largest volume) (Fig. 1F). Collectively, these intravital imaging data reveal that HF-SC and HG progenitor regions have different cell size dynamics *in vivo*. The quiescent bulge resists to cell size changes whereas HGs constantly change cell sizes during both quiescence and activation.

Cell size homeostasis has been associated with cell growth and global protein degradation (12, 18–20). However, the differential size change observed in quiescent HF-SC and HG cells suggested a more dynamic mechanism, which modulates the cell size. We hypothesized that the spatially confined deformation patterns of the bulge and HG could be caused by different mechanical properties, such as different mechanical forces and differential resistance to the forces. We first measured the stiffness of the bulge and HG in the first adult telogen and anagen *ex vivo* by using atomic force microscopy (AFM, fig. S3, A and B). Interestingly, the bulge was 60% stiffer than HGs in the first telogen (1,196Pa vs 747Pa) although these two regions were spatially close. Notably, the stiffness of both bulge and HGs was reduced to 798Pa and 375Pa, respectively, in the first anagen when HF-SCs and HGs were activated to fuel hair growth (Fig. 1, G and H). Because the first telogen is unusually short, typically lasting 3~5 days, we further measured the stiffness of the bulge and HG during the second, prolonged telogen and during the extended telogen in old mice. Both bulge and HGs became progressively stiffer such that the stiffness of the bulge and HG was 2,240Pa and 824Pa, respectively, in P54 samples and 2,620Pa and 1,207Pa, respectively, in 26mo samples (Fig. 1, I and J). The mechanical architecture of the HF-SC compartment, where HF-SCs are stiffer than HG progenitors, is in contrast to that of the epidermis, where differentiated cell layers were >4 times stiffer than the basal SC layer (7).

Next, we examined internal force generation by the bulge and HG cells. Previous studies demonstrated that the actomyosin network plays important roles in pulsed cellular activities during tissue morphogenesis, including apical constriction during gastrulation of *Drosophila* and *C*.*elegans* (21, 22), dorsal closure (23) and germband extension (24) of *Drosophila* and compaction of mouse embryos (25). We quantified filamentous-actin (F-actin) in both bulge HF-SCs and HG cells in telogen and during the telogen-to-anagen transition. To mark the onset of anagen, we used Ki67 staining as an indicator for the active cell cycle. In telogen, HF-SCs located in the bulge had very strong F-actin signals with robust stress fibers across the cells. During early anagen, actin organization transitioned from stress fibers to more cortical organization (Fig. 1K). In contrast to bulge HF-SCs, HG cells exhibited weaker F-actin signals characterized by cortical organization. However, F-actin bundles in HG cells were also rearranged during the activation of anagen hair growth. In telogen, cortical F-actin had stronger signals and were evenly distributed underneath the plasma membrane. At the onset of anagen (anagen I), as Ki67 positive (Ki67+) cells appeared in the HG, F-actin signals became weaker and more diffuse. As the hair cycle progressed to anagen II and more HG cells became Ki67+, HG cells exhibited more diffuse and weaker cortical F-actin signals than their telogen counterparts (Fig. 1K).

We next determined how actomyosin contractility is translated into biochemical signals to activate cell cycle re-entry in the HG. To this end, we examined the correlation between F-actin signal strength and nuclear YAP, a mechanosensitive indicator and signal transducer (5, 26, 27), during telogen and early anagen. YAP signals were weak and diffuse in the cytoplasm during telogen (fig. S4A). Transitioning into early anagen, however, YAP accumulated in the nuclei of a few HG cells (fig. S4B), and the nuclear accumulation spread to nearly all HG cells by late anagen II (fig. S4C). Co-quantification of cortical F-actin and nuclear YAP signals within individual cells revealed an inverse correlation between cortical actomyosin cytoskeleton and nuclear accumulation of YAP (fig. S4D). A randomized test with 1,000 resamplings indicated that the inverse correlation was highly significant (p<0.001) (see Method). Furthermore, a combined F-actin and nuclear YAP score accurately demarcated telogen and anagen states in a mathematical model (fig. S4, E and F). Together, these data reveal an inverse correlation between the actomyosin contractility and nuclear YAP accumulation. At the onset of anagen growth, the weakened actomyosin contractility is correlated with nuclear YAP accumulation in HG progenitors, initiating HG proliferation to fuel hair growth.

### miR-205 induction promotes hair regeneration

We next examined whether the differences in actin cytoskeleton and contractile behaviors observed in bulge HF-SCs and HG cells were associated with differential gene expression. We calculated an actin related gene expression score (see Method) for epithelial and dermal cell populations of the skin by analyzing single-cell RNA-seq datasets from telogen skin (16). HF-SCs located in the bulge had the highest actin score, correlating with the strongest F-actin bundle formation. HG progenitors had lower actin scores than that of HF-SCs (Fig. 2A), consistent with the weaker actomyosin network in the HG. Interestingly, two basal populations of the interfollicular epidermis (IFE), commonly identified in scRNA-seq dataset (28), had different actin scores, indicating that different actin cytoskeleton contributes to epidermal heterogeneity. Sebaceous glands had the lowest actin score among major epithelial populations.

**Figure 2.**
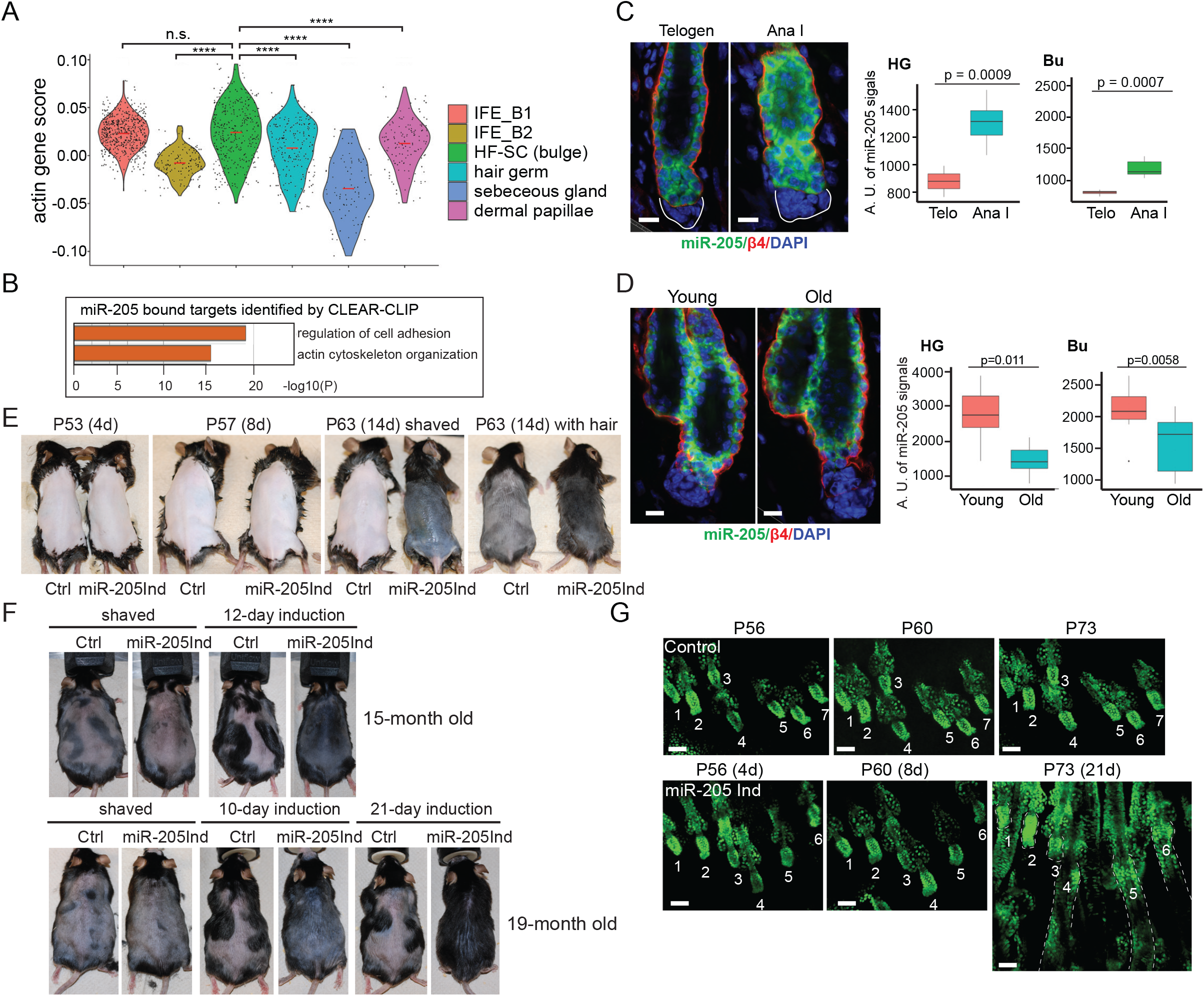
miR-205 induction promotes hair regeneration in young and old mice. **(A)** Epithelial and dermal cell populations from telogen skin had different actin gene expression scores. ****p<0.001, n.s., not significant. IFE_B: interfollicular epidermis basal cell; HF-SC: hair follicle stem cells. **(B)** *miR-205* bound targets were enriched in components and regulators of the actin cytoskeleton and cell adhesion, determined by CLEAR-CLIP. **(C)** In the bulge and HG, the expression of *miR-205* was elevated by ~50% from telogen to the initiation of anagen in the bulge and HG, detected by fluorescent in situ hybridization. Quantification for HG and Bu was showed in the right panel. **(D)** The expression of *miR-205* was decreased by ~100% between young and old telogen HG, and ~ 20% between young and old bulge, detected by fluorescent in situ hybridization. Quantification for HG and bulge was showed in the right panel. (**E)** *miR-205* promoted hair growth in second telogen, starting induction at P49. 8 pairs, 3 pairs and 7 pairs of animals were used for 4-day, 8-day and 14-day induction phenotypical analysis, respectively. **(F)** *miR-205* promoted hair regeneration in old mice (15-m and 19-m old). 4 pairs and 3 pairs of animals were used for 15-month and 19-month induction phenotypical analysis, respectively. **(G)** *miR-205* promoted ear hair follicle regeneration in young mice, observed with a multiphoton microscope. Scale bar 10 μm in **C, D**, 100 μm in **G**. P values were determined by Student’s t-test.

We next asked if we can modulate actomyosin cytoskeleton and force generation by perturbing the expression of actin regulators. We turned to miRNAs, which are known to post-transcriptionally downregulate many genes on a relatively minor scale and in a reversible manner (29, 30). We performed genome-wide capture of miRNA targets together with their associated miRNAs, which detected individual miRNA:mRNA interactions through a direct ligation of miRNA and targeted mRNA site when they were present in the same miRNA-induced silencing complex (31). By analyzing all miRNA-mRNA targeting pairs, we found that *miR-205*, one of the most highly expressed miRNAs expressed in epidermal stem and progenitor cells (32), targets many components and regulators of actomyosin cytoskeleton and cell adhesion. Genes involved in force generation by myosin II activation, such as kinase *Rock2*, and F-actin polymerization, such as *Actb* and *Actn4*, as well as AJ components and force transducers, such as *Ctnna1* (the gene encoding a-catenin), were broadly targeted by this miRNA (Fig. 2B, fig. S5, A and B). Notably, *miR-205* also directly bound to *Piezo1*, a mechanosensitive calcium channel gene, at an evolutionarily conserved site in the 3’UTR (fig. S5B).

We next examined the expression and regulation of *miR-205*. The expression of *miR-205*, determined by quantitative fluorescent in situ hybridization, was elevated by ~50% from telogen to the initiation of anagen in the bulge and HG (Fig. 2C). Furthermore, *miR-205* was downregulated both in the bulge and HG in old mice (Fig. 2D). *miR-205* is transcriptionally regulated by 1 ΔNp63 through three enhancers (fig. S6A), the master transcription factor for specifying epithelial fate of the skin (33). Interestingly, 1 ΔNp63 expression was low and sparse in both HF-SCs and HGs in telogen, compared to their anagen counterparts, while 1 ΔNp63 expression was generally strong and uniform in the IFE basal cells (fig. S6, B to D). In old mice, 1 ΔNp63 expression was significantly reduced in both HF-SCs and HGs, compared to young mice (fig. S6, E to G). Thus, decreased *miR-205* expression was correlated with reduced hair regeneration in old mice.

We next tested whether *miR-205* induction controls bulge HF-SCs and HG cells and modulates hair regeneration. We first induced *miR-205* during early telogen of the second adult hair cycle at P42 (fig. S7, A and B). Usually, the second telogen lasts more than 30 days from P42 to P70 (13). Here, we found that by 5 days after the initial induction (P47), the hair cycle of the dorsal skin transitioned to anagen II-III. By 7 days post induction (P49), the hair cycle reached anagen IV accompanied by the appearance of darkened skin, caused by terminal differentiation of the melanocytes (fig. S7, C and D). The rapid telogen-to-anagen transition caused by *miR-205* was unusual. In comparison, in mice with genetic deletion of *Foxc1*, which is a key transcription factor (TF) governing HF-SC quiescence through the control of BMP and FGF signaling, it takes ~14 days to reach anagen III (34, 35). To further examine the potency of *miR-205* induction on promoting hair growth, we induced *miR-205* at P49, when the dorsal HFs were uniformly in refractory telogen with high BMP signaling (36). By P53, 4 days after the initial *miR-205* induction, both control and induced HFs still appeared to rest in telogen. By P57, 8 days after the initial induction, most dorsal HFs entered anagen III when HF downgrowth was readily observed. By P63 (14 days of induction), the dorsal hair coat was regenerated (Fig. 2E and fig. S7E). During aging, the duration of quiescent telogen gradually increases, and it often lasts >100 days in mice more than 1-year old (37). To test the efficacy of *miR-205* induction in old animals, we examined 15-mo old and 19-mo old mice. In all cases examined, *miR-205* induction resulted in robust and uniform hair growth typically within 14 days, judging by the appearance of darkened skin color and hair coat regeneration (Fig. 2F).

To track the progression of hair growth of individual HFs in live animals, we used 2-photon microscopy to monitor the same HFs (ear) in control and *miR-205* induced animals. Although HFs in the ear were more refractory to the initiation of hair growth (38), *miR-205* induction still initiated anagen hair growth, judging by the HG morphology, within 14 days in young (~P52), middle aged (~12-mo) or older (~15-mo) mice (Fig. 2G and fig. S7F). Thus, *miR-205* induction potently promotes hair regeneration regardless of the age or the inhibitory microenvironment.

### miR-205 inhibits actomyosin contractility and force generation

To determine the underlying mechanism of *miR-205*-mediated hair regeneration, we performed bulk RNA-seq with FACS-purified HF-SCs 2 days after the induction during the refractory telogen before any signs of cell proliferation and anagen re-entry. While *miR-205* was only mildly induced by ~1.8-fold (Fig. 3A), it caused widespread downregulation of targeted genes involved in actin cytoskeleton, cell adhesion, and junctions (fig. S8A). This result confirmed that *miR-205* downregulated genes involved in the regulation of actin cytoskeleton upon the induction in HF-SCs *in vivo*, including *Actb, Actn1, Actn4, Ctnna1, Rock2* and *Piezo1*, identified by the direct ligation of *miR-205* and targeted sites (Fig. 3, B and C and Dataset S1). We next examined the impact of *miR-205* on the transcriptome of epithelial cell populations by performing scRNA-seq 4 days after the induction when HFs transition from telogen to anagen. Unchanged UMAP cell clustering between control and *miR-205* induced samples indicated that *miR-205* induction did not globally perturb cell fate (fig. S8, B to E). Consistent with the bulk RNA-seq data, *miR-205* targets, including the ones involved in myosin II activation, F-actin bundle formation and cell-junction organization, were broadly downregulated in both HF-SCs and HGs (fig. S8, F and G, and Dataset S2). Interestingly, the actin score, a composite index of actin-related gene expression, was significantly downregulated in the actin^hi^ IFE, HF-SCs, HGs and SGs but not in the actin^lo^ IFE and dermal papillae in the *miR-205* induced sample (Fig. 3D), confirming the broad impact of *miR-205* induction on actin cytoskeleton in the epithelial cells. In HF-SC and HG cell clusters, the most highly enriched gene groups among the downregulated *miR-205* targets were actin cytoskeleton and cell adhesion (fig. S8F). Furthermore, genes associated with response to FGF signaling, regulation of mitotic cell cycle and active WNT signaling were enriched among upregulated genes in *miR-205* induced HGs (fig. S8, H and I), consistent with the gene signatures detected during normal telogen-to-anagen transition (13). Together, these data indicate that *miR-205* suppresses genes associated with the actomyosin cytoskeleton and promotes the anagen activation.

**Figure 3.**
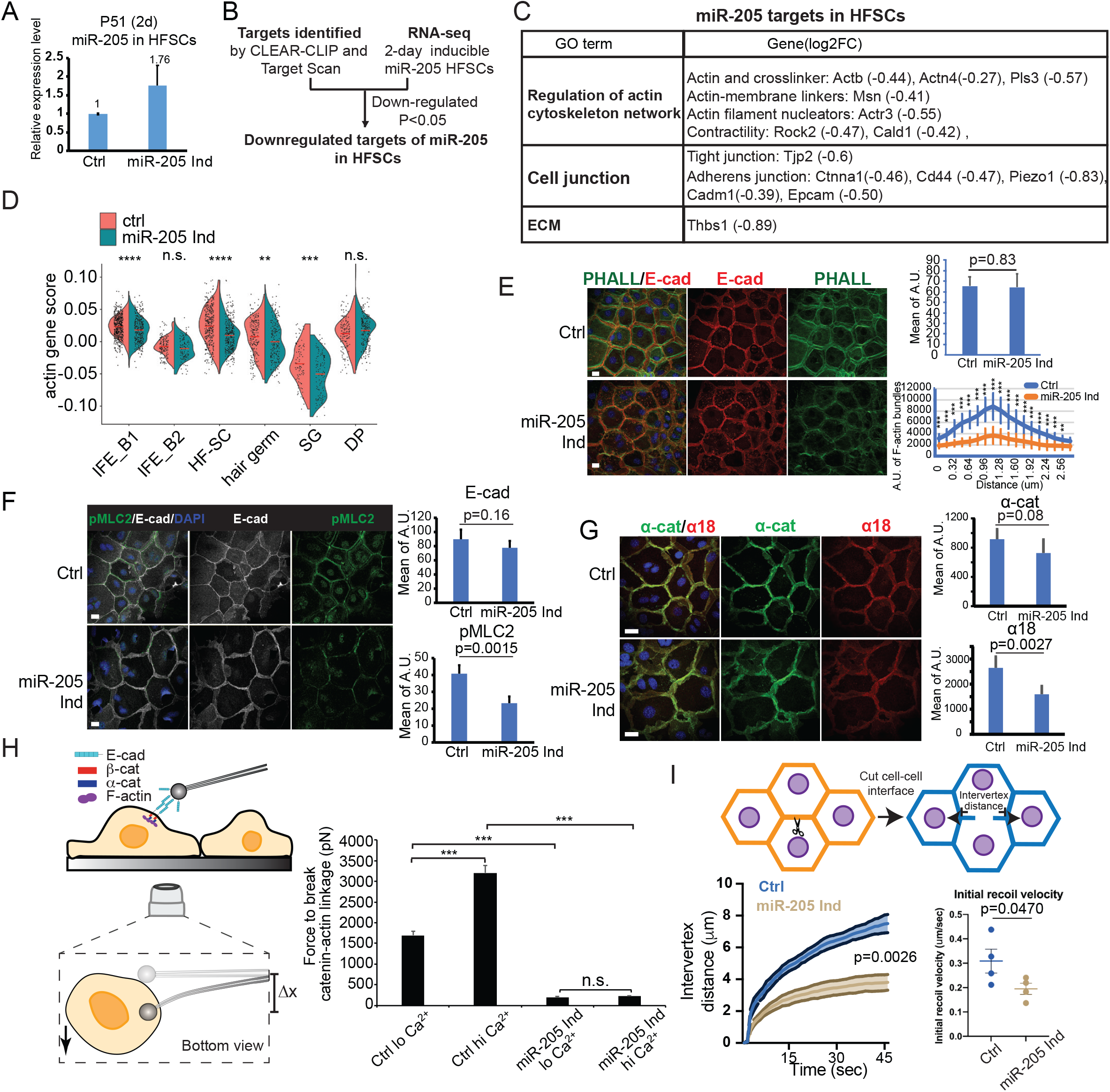
miR-205 targets actomyosin network and inhibits actomyosin contractility and force generation. **(A)** *miR-205* level was increased by ~1.76 fold 2-day after induction in FACS purified HF-SCs. **(B)** Workflow of *miR-205* target identification by combining CLEAR-CLIP capture and TargetScan prediction (for target site conservation), and bulk RNA-seq in HF-SCs. **(C)** miR-205 targets, enriched in the gene categories of cell junction and the actin cytoskeleton, were broadly downregulated in HF-SCs after 2-day induction. **(D)** *miR-205* reduced the actin score in the actin^hi^ basal epidermis, HF-SCs, HGs and SGs but not in DP. (**E)** *miR-205* reduced cortical F-actin bundle formation but not E-cadherin membrane localization. For E-cadherin and PHALL data, 9 control and 15 *miR-205* induction images were used for quantification (upper right). F-actin bundle data (lower right) were mean and s.d. from over 10 line-quantification values. Scale bar, 20 μm. ** P<0.01, *** P<0.001, **** P<0.0001 determined by Student’s t-test. **(F)** *miR-205* reduced pMLC2 levels in keratinocytes. 3 control and 6 *miR-205* induction images were used for quantification (lower right). P values were determined by Student’s t-test. Scale bar, 20 μm. **(G)** *miR-205* reduced actomyosin forces on a-catenin (reflected by a18 signals). 7 control and 4 *miR-205* induced images were used for quantification (right). P values were determined by Student’s t-test. Scale bar, 10 μm. **(H)** *miR-205* reduced the mechanical forces generated by the AJs and the underlying actomyosin networks in keratinocytes with and without calcium induced differentiation. *** P<0.001 determined by Student’s t-test. **(I)** *miR-205* decreased cell membrane tension. For initial recoil velocity assay, P value was determined by Paired t-test. For intervertex distance measurement, P value was determined by Two-way ANOVA. Both graphs showed the average of 4 independent experiments with the error bars as standard error of the mean.

We next examined how elevated *miR-205* expression regulates actomyosin contractility by using primary keratinocyte isolated from the inducible model. Consistent with *in vivo* induction, *miR-205* levels were induced by ~75% 24-hour post induction (fig. 9A). Importantly, induction of *miR-205* reduced cortical F-actin bundle formation without changing the levels of E-Cad, which is not a target (Fig. 3E). Phosphorylated myosin light chain 2 (pMLC2), a well-established effector for actomyosin contractility and regulated by ROCK2 (39–41), was reduced by ~40% in *miR-205* induced keratinocytes (Fig. 3F). Furthermore, whereas the levels of a-catenin (*Ctnna1)*, which is a target of *miR-205*, were mildly reduced (~20%) by *miR-205* as expected, a18 signals were reduced more strongly by ~40% (Fig. 3G), reflecting the reduction of mechanical forces on a-catenin.

To quantify the effect of *miR-205*-mediated regulation on mechanical forces generated by the AJ and the actomyosin cytoskeleton, we measured the forces that were required to break the interaction between AJs and the underlying actomyosin network by using an E-cad microbead displacement experiment (6). In control cells, the forces required to break the AJ-actin linkage were ~1,700 pN in low Ca^2+^ cultured keratinocytes whereas the breaking forces were elevated to ~3,300 pN in high Ca^2+^ cultured, differentiated keratinocytes, reflecting the increased bond between the AJ and the actin cytoskeleton upon Ca^2+^ stimulation (Fig. 3H). Strikingly, the mechanical forces required to break the AJ-actin linkage were reduced by ~10-fold upon *miR-205* induction, regardless of the Ca^2+^ concentration in the media (Fig. 3H). In comparison, genetic deletion of *VCL*, a key connector of AJs to the actin cytoskeleton, reduced the mechanical forces by ~2-fold measured by the same assay (6). In mice, genetic deletion of *VCL* strongly reduced F-actin and a18 signals and caused widespread cell cycle re-entry of both HF-SCs and HGs (fig. S9, B and C), directly linking AJ-mediated force generation to the control of cell cycle re-entry. Collectively, these results demonstrate that co-targeting of myosin II activation, F-actin bundle formation and AJ components by *miR-205* substantially reduces the AJ-actin force generation.

Finally, we asked whether *miR-205* induction also perturbed the tension on the plasma membrane because membrane tension could constrain cell size. A laser ablation and recoil assay (42, 43) revealed that the intervertex distance was reduced by ~50% in 45 seconds after the laser ablation and the initial recoil velocity was reduced by ~33% (Fig. 3I). Together, these data demonstrate that *miR-205* induction reduces mechanical forces generated by the AJs and actomyosin contractility at the cortex and decreases the membrane tension.

### Mechanism of miR-205-mediated hair regeneration

Having established the role of *miR-205* in modulating actomyosin contractility and membrane tension *in vitro*, we examined cellular responses to *miR-205* induction *in vivo*. The appearance of Ki67+ cells in the HG marks the onset of anagen initiation. In *miR-205* induced skin, Ki67+ HG cells began to emerge sporadically 4 days after the initial induction (P49-P53), and ~20% of HFs contained Ki67+ HG cells. By 5 days after the induction, however, nearly 100% of HFs contained Ki67+ HG cells (Fig. 4, A and B). These data were consistent with our transcriptome analysis for the effect of *miR-205* induction. Indeed, cortical F-actin bundles were reduced in HG cells 4 days after the induction (Fig. 4C, left panels). By 5 days after the induction, cortical F-actin signals were further weakened and became diffuse with clear signs of cell number increase in the HG (Fig. 4C, right panels), reflecting cell cycle re-entry. Consistent with these morphological changes in actomyosin networks, mechanical properties of HG cells also changed, indicated by reduced (~40%) and punctate a18 signals that reflect weakened actomyosin forces on a-catenin (Fig. 4D). The reduced a18 signals in *miR-205* induced HGs were similar to those of the normal telogen-to-anagen transition, observed in control mice (fig. S9D). Accompanying these changes, YAP protein began to accumulate in the nuclei of not only HG cells but also bulge HF-SCs 4-day after *miR-205* induction (Fig. 4E). Furthermore, modeling of cortical F-actin and nuclear YAP signals obtained from *miR-205* induced samples revealed that *miR-205* induction promoted the transition of HG cells from the quiescent telogen state to the activated anagen state (fig. S9, E and F). In addition to the changes in actin cytoskeleton, *miR-205* also reduced the stiffness of both the bulge and HG compartments in both young and old samples, determined by AFM ex vivo (Fig. 4, F to I). In P54 samples, bulge stiffness was reduced from 2,240Pa to 2,030Pa, and HG stiffness was reduced from 824Pa to 505Pa (Fig. 4, F and G). Consistent with the potent induction of hair regeneration by *miR-205*, the stiffness of the bulge (2,620Pa) and HG (1,207Pa) in 21-22mo old mice was reduced to the levels usually found in young mice (bulge, 2,335Pa; HG, 887Pa) (Fig. 4, H and I).

**Figure 4.**
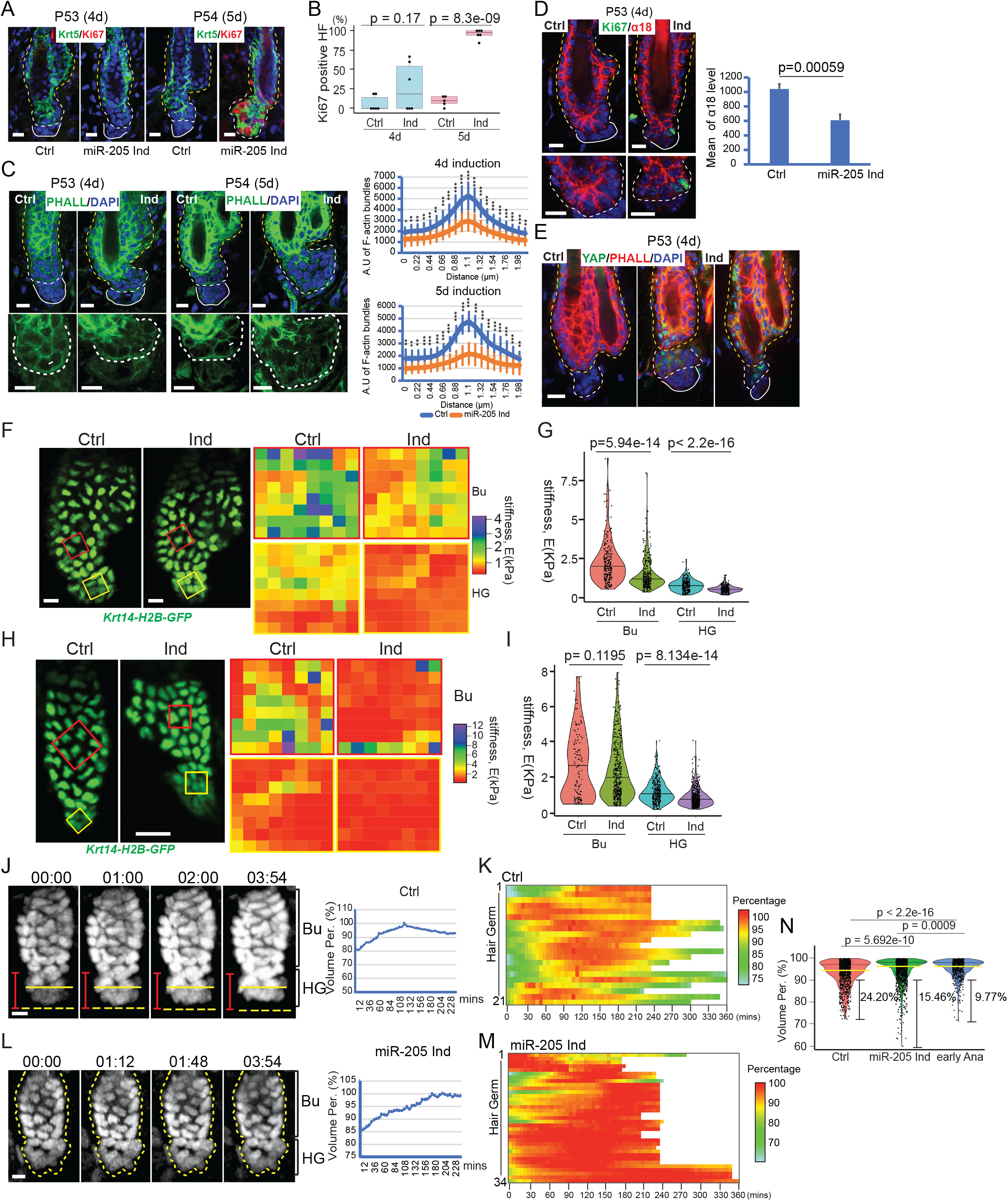
miR-205 modulates mechanical stiffness and actomyosin contractility of HF-SC compartment. **(A)** *miR-205* promoted Ki67 accumulation 4-5 days after induction. **(B)** The percentages of Ki67+ HFs were quantified after 4-day and 5-day induction. 71 control HFs and 57 *miR-205* induced HFs from 6 pairs of animals for 4-day induction, 61 control HFs from 5 animals and 145 *miR-205* induced HFs from 6 animals for 5-day induction, respectively, were used. **(C)** *miR-205* reduced cortical F-actin levels and modulated the actomyosin network after 4-day and 5-day induction. F-actin bundle data (right panels) were mean and s.d. from over 10 line-quantification values. **(D)** *miR-205* reduced actomyosin forces on a-catenin (reflected by a18 signals). 7 control and 4 *miR-205* induced images were used for quantification (right). P values were determined by Student’s t-test. **(E)** *miR-205* induction remodeled the actomyosin cytoskeleton and resulted in nuclear YAP accumulation in both HF-SCs and HG cells after 4-days induction. **(F)** *miR-205* reduced the stiffness of both the bulge and HG compartments after 5-day induction in young mice, determined by AFM ex vivo. **(G)** The stiffness of both the bulge and HG compartments was reduced in *miR-205* induced old hair follicles. Bu in control skin, mean= 2240.1 ± 1251.1 Pa; Bu in miR-205 samples, mean= 2030.8 ± 1570.6 Pa. HG in control samples, mean= 824.2 ± 368.0 Pa; HG in miR-205 skin, mean= 505.7 ± 244.6 Pa. **(H)** *miR-205* reduced the stiffness of both the bulge and HG compartments after 7-day induction in old mice, determined by AFM ex vivo. **(I)** The stiffness of both the bulge and HG compartments was reduced in *miR-205* induced hair follicles in old mice. Bu in control skin, mean= 2620.9 ± 1868.0 Pa; Bu in miR-205 samples, mean= 2335.6 ± 1603.0Pa. HG in control samples, mean= 1207.2 ± 625.2 Pa; HG in miR-205 skin, mean= 886.9 ± 501.9 Pa. **(J)** An example of HG undergoing enlargement and contraction (left) with quantification (right) over 4 hours in second telogen. **(K)** HGs showed the cyclic behaviors of contraction and enlargement in second telogen. 21 hair follicles in second telogen from 7 control animals were quantified. **(L)** An example of HG undergoing continuous enlargement (left) with quantification (right) in *miR-205* induced sample over 4 hours. **(M)** *miR-205* reduced HG contraction. 34 hair follicles from 8 *miR-205* induced animals were recorded. (**N)** miR-205 induced HGs reduced contraction, mimicking the behavior observed in early anagen of control mice. The yellow lines represented the median values and the black lines represented the third quarter of all time points in the violin plot. Scale bar 10 μm in **A, C, D, E, F, J, L**, 20 μm in **H**. P values were determined by Student’s t-test.

Because *miR-205* changed mechanical properties by reducing actomyosin contractility and the stiffness of the bulge and HG, we next asked whether *miR-205* changes contractile behaviors and cell size dynamics of HGs in live animals. During the second, prolonged telogen, time-lapse live imaging revealed the same enlargement and contraction activities of quiescent HGs. In 21 HFs from 7 mice, 19 (90.5%) HGs exhibited dynamic enlargement and contraction with a size change of ~20% and 2 (9.5%) HGs only underwent enlargement during the 4~6-hour imaging window (Fig. 4, J and K). These cell behaviors were reminiscent to the behavior of HGs in the first telogen, during which 93.5% of HGs (29 out of 31 HFs) exhibited substantial (>10%) contraction (see Fig. 1B). In contrast, only 9 (26.5%) HGs showed substantial (>10%) contraction whereas the remaining 73.5% of HGs did not contract or mildly increased their size in 34 HFs from 8 induced mice (Fig. 4, L and M and movie S5). Thus, *miR-205* induction significantly reduced HG contraction, defined by the percentage of HGs whose size was less than 90% of the largest volume during observation (Fig. 4N). Collectively, these results demonstrate that *miR-205* induction reduces cell size contraction and promotes cell cycle re-entry through reduced actomyosin contractility and stiffness in the bulge and HG.

## Discussion

In conclusion, we have determined spatiotemporal compartmentalization of mechanical properties, including stiffness and actomyosin contractility, and demonstrated their impacts on cell size and cell activities within the HF-SC niche in live animals (fig. S9G). The high stiffness and strong actomyosin contractile forces of the bulge impose tight constraints on cell size changes, and they collectively contribute to the quiescence control of bulge HF-SCs in homeostasis and during aging. In contrast, HG progenitors, which exhibit lower stiffness and weaker actomyosin contractility, are soft and mechanically active. Cell size control and cell mechanics are fundamental to cell physiology and tissue homeostasis (1, 2, 12, 18, 44). This study has implicated the dynamic changes of cell size and subsequent cell cycle re-entry as a mechanism to sense and respond to mechanical forces.

## Materials and Methods

### Cell lines

Primary keratinocytes were isolated from mice between the ages of P0 and P4 as previously described, with some modifications(45). Mice were sacrificed, back and belly skin was collected, and excess fat was scraped from the tissue. The skin was then placed dermal-side down in 2× dispase for 30–60 min at 37°C. The epidermis was subsequently separated from the dermis with forceps and placed into 0.05% trypsin-EDTA for 10 min at 37°C. Trypsin was quenched with culture media, and cells were strained through a 40-μm filter and subsequently plated in E-low Ca2+ media onto dermal feeder cells.

### Animal studies

All mice breeding and operation procedures were approved by the institutional animal care and use committees (IACUC) at University of Colorado Boulder (CO, USA) and at Northwestern University Feinberg School of Medicine (IL, USA), and in accordance with the guidelines and regulations for the care and use of laboratory animals.

The miR-205 induction mouse line was generated through standard transgenic injection of the linearized *pTRE2-miR-205* DNA into FVB mice. Founders were bred with mice harboring a keratin14 reverse tetracycline trans-activator (*Krt14*-rtTA, E. Fuchs, Rockefeller University) to produce mice with skin-specific doxycycline-inducible expression of the miR-205. Multiple lines were established and validated for the study. Doxycycline chow used in experiments was 625 mg/kg (Teklad rodent diet TD-7012). E-cadherin-mCFP (JAX # 016933), Rosa-mTmG (JAX # 007676) and Rosa26-LSL-tdTomato (JAX # 021876) mice were obtained from the Jackson laboratory. *Krt14*-*H2b*-*GFP* (E. Fuchs, Rockefeller University), E-cadherin-mCFP, Rosa-mTmG and Rosa26-LSL-tdTomato mice were used for live animal imaging.

### Immunostaining, in situ hybridization, and imaging

For analysis of back skin phenotypes, 10 μm OCT sections were fixed in 4% PFA for 10 min in phosphate buffered saline (PBS) and washed three times for 5 min in 1x PBS at room temperature. Block the sections with 2.5% NGS, 2.5% NDS in PBS. Sections were incubated with primary antibody overnight at 4°C. After incubation with primary antibodies, sections were washed three times in 1× PBS and incubated for 1 h at room temperature with Alexa Fluor 594–, Alexa Fluor 488–, or Alexa Fluor 647–conjugated secondary antibodies (1:2,000; Invitrogen/Molecular Probes). Alexa Fluor™ Plus 555 Phalloidin (1:200; A30106; Invitrogen) or Alexa Fluor 488 conjugated phalloidin (1:50; A12379; Invitrogen) were used to stain F-actin. Nuclei were stained with Hoechst 33342 (1:5000; Invitrogen). Imaging was performed on a Leica DM5500B microscope with an attached Hamamatsu C10600-10B camera and MetaMorph (version 7.7; MDS Analytical Technologies) software, and Nikon W1 Dual CAM Spinning Disk at Northwestern University Feinberg School of Medicine.

In situ hybridizations were performed by adapting miRNA localization protocols to skin. Signals were detected using the TSA Plus Fluorescein System (Perkin Elmer) for fluorescent signal and post-hybridization staining. The miRCURY LNA miR-205 detection probe (Exiqon) was 5’-DIG and 3’-DIG double labelled and hybridized to skin sections at 61 °C. Sections were stained with β4-integrin (BD Biosciences, 553745) and Alexa-Fluor-594-conjugated secondary antibody (1:2,000, Invitrogen/Molecular Probes, A11007) after developing.

### Fluorescence-activated cell sorting (FACS)

Gender matched mice (Ctrl and miR-205 Ind) harboring a *Krt14*-*H2b*-*GFP* allele were fed doxycycline for 2 or 4 days and euthanized to collect back skin samples. Hair coats were shaved and removed by nair hair removal lotion (Amazon, 22339) for around 5 minutes. Subcutaneous fat was removed, and back skin samples were minced into small pieces and incubated with 0.25% collagenase (Worthington, LS004188) in 10 mL 1x HBSS buffer at 37°C for 2 hours with rotation. 25 mL serological pipet was used to further enhance collagenase reaction at 1 hour incubation time. After collagenase treatment, 10 mL cold PBS was added, and sample suspension was centrifugated at 400 g for 10 minutes at 4°C to get the pellet. The pellet was resuspended with 5 ml pre-warmed 0.25% trypsin/EDTA (Gibco) for 7 minutes at 37°C and the digestion was blocked by adding 10 mL cold 1x PBS with 3% chelated FBS. The suspension was extensively triturated with a 25 ml pipette and filtered through a 40 μm cell strainer, followed by centrifugation at 400 g for 5 minutes at 4°C. The pelleted cells were resuspended in cold 1x PBS with 3% chelated FBS. Suspended cells were incubated with appropriated antibodies for 1 hour on ice. After washing away unbounded antibodies, DAPI was used to exclude dead cells. Hair follicle stem cells (HF-SCs) of Krt14-H2b-GFP-based experiments were isolated by enriching DAPI^neg^, H2BGFP^hi^, Sca1^neg^, α6^hi^ and CD34^hi^ cells. The following antibodies were used in K14-H2b-GFP-based experiments: integrin α6 (CD49f, 1:50; eBioscience, PE-conjugated, 12-0495), CD34 (1:100; eBioscience, eFluor 660-conjugated, 50-0341), Sca1 (Ly-6A/E, 1:500; eBioscience, PE-Cyanine7-conjugated, 25-5981-82). FACS was performed on MoFlo XDP machine (Beckman Coulter). FACS data were analyzed with FlowJo.

### RNA Purification and miRNA qPCR

Total RNA was extracted using TRIZOL (Thermo Fisher Scientific). For miRNA analysis, miScript II RT Kit (Qiagen) was used to synthesize cDNA. miScript primers for miRNA-205 and control Sno25 were from Qiagen. Reactions were performed according to the manufacturer’s manual and on a CFX384 real-time system (Bio-Rad). Differences between samples and controls were calculated using the 2^-ΔC(t)^ method.

### Laser ablation and recoil assay

CellMask™ Orange Plasma membrane Stain (ThermoFisher, C10045) was used to visualize mouse keratinocytes according to manufacturers’ instructions. Mouse keratinocyte cultures were allowed to equilibrate for 20 minutes before imaging and only imaged for 1 hour each. Two-photon laser ablation was used to assess intercellular forces. Briefly, ablation was performed on a Nikon A1R-MP+ multiphoton microscope running Elements version 4.50 and equipped with an Apo LWD 25× 1.10W objective. Cells were maintained at 37 °C and 5% CO2. Images were obtained at a rate of 1 frame per second for 2 s before and 45 s after ablation using 4% laser power at 970 nm. Ablation was performed using 40% laser power at a scan speed of 512. The distance between the cell–cell vertices over time was measured using the Manual Tracking plugin in ImageJ. Distance curves were then generated using Excel software. Initial recoil measurements were calculated using Prism 8 software as previously described(46).

### Force measurements at the AJs

The microbeads (Dynabeads M270, Invitrogen) were treated by 1% glutaraldehyde for 1 hour at room temperature. After washed with 1 X PBS for five times, the beads were incubated in 10 μg/ml recombinant E-cadherin (human, Sigma) solution overnight at 4 °C with agitation. The beads were then washed and stored in blocking buffer (1% BSA, 1 X PBS) at 4 °C on an agitation rotator for use.

Keratinocytes were seeded in a home-made chamber coated with 10 μg ml^-1^ fibronectin. For the miR-205 inducible cells, cells were treated with 3 μg/ml final concentration doxycycline after cell attachment. For high Calcium treatment, 2μM final concentration Calcium was applied. After 24 hours incubation, the E-cadherin coated microbeads were added into the chamber. The glass micropipette mounted on a MP-285 micromanipulator was gently inserted into the chamber with a small angle, and captured an E-cadherin coated bead through liquid aspiration pressure. Then the bead was moved down to touch the cell surface for 1 minutes to allow E-cadherin mediated AJ formation. The cell was subsequently moved away from the micropipette by controlling the microscopy stage at a speed of 10 μm s^-1^, resulting in the micropipette bending until the adhesion interface rupture (Figure 4E). The force (F) at which the bead was detached from the cell surface was a direct readout of the strength of the E-cadherin mediated AJs and underlying actomyosin formed by these cells with the beads. This helped to quantify the contribution of α-catenin and underlying actomyosin to the adhesion strength at the AJs.

The micropipette experiments were recorded and analyzed by ImageJ to determine the micropipette deflection distance (Δx). The amount of force can be evaluated from the micropipette deflection distance (Δx) based on the pre-calibrated bending stiffness (k) of the micropipette (F=kΔx).

### RNA-seq assay

Total RNAs from FACS-purified cells were isolated using TRIZOL (Invitrogen). RNA quality was assessed by Agilent 2100 bioanalyzer. RNA Samples with RNA integrity numbers (RIN) > 8 were used to make libraries and perform RNA-seq assay. Libraries were prepared by following manufacturer’s protocol (NEBNext Ultra Directional RNA Library Prep Kit). The cDNA libraries were quality-checked with bioanalyzer and sequenced at Genomics and Microarray Core Facility at University of Colorado Denver on Illumina NovaSeq 6000.

### RNA-seq analysis

RNA-seq reads (150nt, paired-end) were aligned to the mouse genome (NCBI37/mm10) using Hisat2 (version 2.1.0)(47). The resulting sam files were converted to bam files using SAMtools (version 1.3.1)(48). Expression measurement of each gene was calculated from the resulting alignment bam file by HT-seq7(49). Differentially expressed genes were determined using DEseq2(50) using default settings, with adjusted p-value cutoff of 0.05. Gene ontology analysis was performed using Metascape(51). The selected GO terms were from the metascape results along with the gene lists.

### Single-cell RNA-seq assay and data analysis

Single-cell RNA-seq was performed according to manufacturer’s instruction (10X Genomics). Cells from both control and miR-205 inducible animals after 4-day induction were collected from FACS sorting machine with cell surface proteins and H2b-GFP signals such that the epidermal cells, hair follicle cells and dermal cells are 3: 5: 2 ratio. Each sample was targeted to obtain ~5,000 cells. Single-cell RNA-seq reads were sequenced on Illumina Hi-Seq 4000.

The Cell Ranger Single-Cell Software Suite was used to perform barcode processing and single-cell 3’ gene counting (https://support.10xgenomics.com/single-cell-gene-expression/software/overview/welcome).The barcodes, features and matrix files were loaded into Seurat 3.0 for downstream analysis (https://satijalab.org/seurat/articles/install.html). For each sample, the analysis pipeline followed the guided tutorial. The cells were filtered with nFeature_RNA (>200 and <5000) and mitochondrial percentage (< 10). After clustering and umap dimension reduction, the cluster markers were used to identify distinct cell populations. For comparison among different samples, samples were integrated to promote the identification of common cell types and enable comparative analysis.

For bulk-RNAseq analysis for outer bulge stem cell (OB) and HG clusters, mapped transcriptome datasets from OB and HG clusters were extracted according to their cell barcodes, combined together and obtained sam files. The sam files were further analyzed as bulk-RNAseq datasets. The resulting sam files were converted to bam files using SAMtools (version 1.3.1)(48). Expression measurement of each gene was calculated from the resulting alignment bam file by HT-seq7(49). Differentially expressed genes were determined using DEseq2(50) using default settings, with adjusted p-value cutoff of 0.05. Gene ontology analysis was performed using Metascape(51). The selected GO terms were from the metascape results along with the gene lists.

### Intravital live imaging and image processing and quantification

Intravital live imaging was performed as previously described(15, 52) with modifications. Mice used for imaging were sedated using 1% oxygen and ~2% isoflurane. Once a mouse was fully sedated, it was put on a warm pad at 37°C. The oxygen and isoflurane were maintained during the course of imaging. Night-time ointment (Genteal, NDC 0078-0473-97) was applied to keep eyes moisturized. Custom manufactured spatula was used to flatten the region of interest and maintained at adjustable height. Double sided tapes were used to adhere the lower side of ear onto the spatula. After applying long-lasting Genteal gel (0078-0429-47) to the region of interest, a second adjustable spatula, glued with cover-glass on one end, was gently pressed down to the ear so that the cover glass was right on top of the region. Long-lasting Genteal was applied on the cover-glass to keep the tip of objective merged in during imaging. Olympus FVMPE-RS multiphoton imaging system was applied for acquiring Krt14-H2b-GFP and Rosa26-LSL-Tdtomato images. The lasers were Insight X3 with wavelength set to 920 nm for GFP signals and 1050 nm for Tdtomato signals. Emission wavelength were 510 nm and 580 nm respectively. 10X and 25X objectives were used for images. The Leica DiveB Sp8 Multiphoton imaging system was used for E-cad-mCFP and mTmG/Krt14-H2B-GFP images. During the imaging session, the light should be turned off and the stages and scope should be covered with black curtain to avoid exposure to light.

After the image session is done, the mouse was kept in oxygen to recover before sending back in cage. Two photon images were acquired using Fluoview software from Olympus. The time lapse images were aligned using Fiji > plugins > registration > descriptor-based series registration (2d/3d +t) before exported. Then images were exported to tif format using Fiji > plugins > bio-formats > bioformats-exporter. The exported tif files were further converted to Imaris file format using Imaris File Converter software. The Imaris x64 9.7.2 were used for further analysis. The images were adjusted on x, y or z plane for better visualization. Movies were also adjusted and generated from Imaris.

For hair germ volume quantifications, the 3D pictures were opened in Imaris and then cut off the epidermis and upper hair follicle regions to acquire individual hair follicles, including bulge and HG. Individual HG regions were applied with Imaris Surface module. The Volume output of hair germ was used for quantification. The hair follicles with obvious drift were excluded to avoid the artificial effects. Individual HG volumes are heterogenous. To determine the dynamic changes of HGs at different stages and in different samples, we designated the largest value of the volume during a period of time as 100% and obtained relative percentage value at different time points by comparing with the largest value. The average cell volumes were obtained from total HG volume divided by cell numbers included for HG volume calculation.

### Actin Score Calculation

Actin Scores were calculated using AddModuleScore function in the Seurat package. We defined a custom actin cytoskeleton organization gene list from GO: GO0030036 by excluding genes that were not detected by our single-cell RNA-seq and genes that we did not find appropriate synonym with geneSynonym package.

### Statistics and study design

In general, all sequencing experiments (RNA-seq) were repeated on at least two pairs of control and miR205 inducible per experiment. Single-cell ATAC-seq was performed with one pair of control and miR-205 inducible samples at the same time on the same chip to avoid the batch effect. All experiments were designed such that there were always littermate controls. All statistical tests performed are as indicated in the figure legends. No statistical methods were used to predetermine sample size. The experiments were not randomized, and the investigators were not blinded to allocation during experiments and outcome assessment, except where stated.

For all experiments with error bars, the standard deviation (s.d.) was calculated to indicate the variation within each experiment. Numbers of animals used for phenotype study has indicated in the manuscript and figure legends. Student’s t-test was used for most experiments.

## Acknowledgments

We thank E. Fuchs (Rockefeller University, HHMI) for Krt14-rtTA, Krt14-H2bGFP and Krt14-Cre mice; A. Nagafuchi (Nara Medical University) for a18 monoclonal antibody; J. Orth (University of Colorado Boulder), J. Lopez and R. Goldmeyer (Olympus) and C. Arvanitis and H. Chen (Northwestern University CAM) for two-photon imaging; D. Klrchenbuechler (Northwestern University CAM) for AFM; and all members of the Yi laboratory for suggestions. This work was supported by National Institute of Health Grant AR066703 and AR071435 (RY), National Science Foundation CBET 2029559 (RY); National Institute of Health Grants AR043380 and AR041836 (KG); National Institute of Health K01 AR075087 (JAB); Singapore Ministry of Education and the National University of Singapore under the Research Scholarship Block (JY).

## References

1. K. H. Vining, D. J. Mooney, Mechanical forces direct stem cell behaviour in development and regeneration. Nat. Rev. Mol. Cell Biol. 18, 728–742 (2017).

2. D. A. Fletcher, R. D. Mullins, Cell mechanics and the cytoskeleton. Nature 463, 485–492 (2010).

3. A. J. Engler, S. Sen, H. L. Sweeney, D. E. Discher, Matrix Elasticity Directs Stem Cell Lineage Specification. Cell 126, 677–689 (2006).

4. L. He, G. Si, J. Huang, A. D. T. Samuel, N. Perrimon, Mechanical regulation of stem-cell differentiation by the stretch-activated Piezo channel. Nature 555, 103–106 (2018).

5. M. Aragona, et al., Mechanisms of stretch-mediated skin expansion at single-cell resolution. Nature 584, 268–273 (2020).

6. R. Biswas, et al., Mechanical instability of adherens junctions overrides intrinsic quiescence of hair follicle stem cells. Dev. Cell 56, 761–780.e7 (2021).

7. V. F. Fiore, et al., Mechanics of a multilayer epithelium instruct tumour architecture and function. Nature 585, 433–439 (2020).

8. J. Koester, et al., Niche stiffening compromises hair follicle stem cell potential during ageing by reducing bivalent promoter accessibility. Nat. Cell Biol. 23, 771–781 (2021).

9. Y. A. Miroshnikova, et al., Adhesion forces and cortical tension couple cell proliferation and differentiation to drive epidermal stratification. Nat. Cell Biol. 20, 69–80 (2018).

10. W. Ning, A. Muroyama, H. Li, T. Lechler, Differentiated Daughter Cells Regulate Stem Cell Proliferation and Fate through Intra-tissue Tension. Cell Stem Cell 28, 436–452.e5 (2021).

11. Y. Barrandon, H. Green, Cell size as a determinant of the clone-forming ability of human keratinocytes. Proc. Natl. Acad. Sci. 82, 5390–5394 (1985).

12. S. Xie, J. M. Skotheim, A G1 Sizer Coordinates Growth and Division in the Mouse Epidermis. Curr. Biol. 30, 916–924.e2 (2020).

13. V. Greco, et al., A two-step mechanism for stem cell activation during hair regeneration. Cell Stem Cell 4, 155–69 (2009).

14. Y.-C. Hsu, L. Li, E. Fuchs, Transit-amplifying cells orchestrate stem cell activity and tissue regeneration. Cell 157, 935–949 (2014).

15. C. M. Pineda, et al., Intravital imaging of hair follicle regeneration in the mouse. Nat. Protoc. 10, 1116–1130 (2015).

16. C. Zhang, et al., Escape of hair follicle stem cells causes stem cell exhaustion during aging. Nat. Aging 1, 889–903 (2021).

17. K. R. Mesa, et al., Homeostatic Epidermal Stem Cell Self-Renewal Is Driven by Local Differentiation. Cell Stem Cell 23, 677–686.e4 (2018).

18. M. B. Ginzberg, R. Kafri, M. Kirschner, Cell biology. On being the right (cell) size. Science 348, 1245075 (2015).

19. R. Kafri, et al., Dynamics extracted from fixed cells reveal feedback linking cell growth to cell cycle. Nature 494, 480–483 (2013).

20. S. Liu, et al., Large cells activate global protein degradation to maintain cell size homeostasis. 2021.11.09.467936 (2021).

21. A. C. Martin, M. Kaschube, E. F. Wieschaus, Pulsed contractions of an actin–myosin network drive apical constriction. Nature 457, 495–499 (2009).

22. M. Roh-Johnson, et al., Triggering a Cell Shape Change by Exploiting Preexisting Actomyosin Contractions. Science 335, 1232–1235 (2012).

23. J. Solon, A. Kaya-Äopur, J. Colombelli, D. Brunner, Pulsed Forces Timed by a Ratchet-like Mechanism Drive Directed Tissue Movement during Dorsal Closure. Cell 137, 1331–1342 (2009).

24. A. Munjal, J.-M. Philippe, E. Munro, T. Lecuit, A self-organized biomechanical network drives shape changes during tissue morphogenesis. Nature 524, 351–355 (2015).

25. J.-L. Maître, R. Niwayama, H. Turlier, F. Nédélec, T. Hiiragi, Pulsatile cell-autonomous contractility drives compaction in the mouse embryo. Nat. Cell Biol. 17, 849–855 (2015).

26. S. Dupont, et al., Role of YAP/TAZ in mechanotransduction. Nature 474, 179–183 (2011).

27. F.-X. Yu, B. Zhao, K.-L. Guan, Hippo Pathway in Organ Size Control, Tissue Homeostasis, and Cancer. Cell 163, 811–828 (2015).

28. S. Joost, et al., Single-Cell Transcriptomics Reveals that Differentiation and Spatial Signatures Shape Epidermal and Hair Follicle Heterogeneity. Cell Syst. 3, 221–237.e9 (2016).

29. V. Ambros, The functions of animal microRNAs. Nature 431, 350–355 (2004).

30. D. P. Bartel, MicroRNAs: target recognition and regulatory functions. Cell 136, 215–233 (2009).

31. G. A. Bjerke, R. Yi, Integrated analysis of directly captured microRNA targets reveals the impact of microRNAs on mammalian transcriptome. RNA 26, 306–323 (2020).

32. D. Wang, et al., MicroRNA-205 controls neonatal expansion of skin stem cells by modulating the PI(3)K pathway. Nat. Cell Biol. 15, 1153–1163 (2013).

33. X. Fan, et al., Single Cell and Open Chromatin Analysis Reveals Molecular Origin of Epidermal Cells of the Skin. Dev. Cell 47, 21–37.e5 (2018).

34. K. Lay, T. Kume, E. Fuchs, FOXC1 maintains the hair follicle stem cell niche and governs stem cell quiescence to preserve long-term tissue-regenerating potential. Proc. Natl. Acad. Sci. U. S. A. 113, E1506–1515 (2016).

35. L. Wang, J. A. Siegenthaler, R. D. Dowell, R. Yi, Foxc1 reinforces quiescence in self-renewing hair follicle stem cells. Science 351, 613–617 (2016).

36. M. V. Plikus, et al., Cyclic dermal BMP signalling regulates stem cell activation during hair regeneration. Nature 451, 340–344 (2008).

37. C.-C. Chen, et al., Regenerative hair waves in aging mice and extra-follicular modulators follistatin, dkk1, and sfrp4. J. Invest. Dermatol. 134, 2086–2096 (2014).

38. Q. Wang, et al., A multi-scale model for hair follicles reveals heterogeneous domains driving rapid spatiotemporal hair growth patterning. eLife 6, e22772 (2017).

39. K. A. Newell-Litwa, et al., ROCK1 and 2 differentially regulate actomyosin organization to drive cell and synaptic polarity. J. Cell Biol. 210, 225–242 (2015).

40. K. Kimura, et al., Regulation of myosin phosphatase by Rho and Rho-associated kinase (Rhokinase). Science 273, 245–248 (1996).

41. M. S. Samuel, et al., Actomyosin-mediated cellular tension drives increased tissue stiffness and β-catenin activation to induce interfollicular epidermal hyperplasia and tumor growth. Cancer Cell 19, 776–791 (2011).

42. O. Nekrasova, et al., Desmosomal cadherin association with Tctex-1 and cortactin-Arp2/3 drives perijunctional actin polymerization to promote keratinocyte delamination. Nat. Commun. 9, 1053 (2018).

43. J. A. Broussard, J. L. Koetsier, M. Hegazy, K. J. Green, Desmosomes polarize and integrate chemical and mechanical signaling to govern epidermal tissue form and function. Curr. Biol. 31, 3275–3291.e5 (2021).

44. D. E. Discher, P. Janmey, Y. Wang, Tissue Cells Feel and Respond to the Stiffness of Their Substrate. Science 310, 1139–1143 (2005).

45. R. Yi, M. N. Poy, M. Stoffel, E. Fuchs, A skin microRNA promotes differentiation by repressing ‘stemness.’ Nature 452, 225–229 (2008).

46. X. Liang, M. Michael, G. A. Gomez, Measurement of Mechanical Tension at Cell-cell Junctions Using Two-photon Laser Ablation. Bio-Protoc. 6, e2068 (2016).

47. D. Kim, B. Langmead, S. L. Salzberg, HISAT: a fast spliced aligner with low memory requirements. Nat. Methods 12, 357–360 (2015).

48. H. Li, et al., The Sequence Alignment/Map format and SAMtools. Bioinformatics 25, 2078–2079 (2009).

49. S. Anders, P. T. Pyl, W. Huber, HTSeq—a Python framework to work with high-throughput sequencing data. Bioinformatics 31, 166–169 (2015).

50. M. I. Love, W. Huber, S. Anders, Moderated estimation of fold change and dispersion for RNA-seq data with DESeq2. Genome Biol. 15, 550 (2014).

51. Y. Zhou, et al., Metascape provides a biologist-oriented resource for the analysis of systems-level datasets. Nat. Commun. 10, 1523 (2019).

52. P. Rompolas, et al., Live imaging of stem cell and progeny behaviour in physiological hair-follicle regeneration. Nature 487, 496–499 (2012).

