## Supplementary_data for "MicroRNA-205 promotes hair regeneration by modulating mechanical properties of hair follicle stem cells"

**This PDF file includes:**
Supporting text
Figures S1 to S9
Legends for Movies S1 to S5
Legends for Datasets S1 to S2
SI References

**Other supporting materials for this manuscript include the following**:
Movies S1 to S5
Datasets S1 to S2

**Supporting Information Text**

**Method**

Atomic force microscopy

25-μm-thick OCT cryosections of back skin for control and miR-205 inducible were prepared for measurements of hair follicles and dermal tissue stiffness. Samples were affixed to Superfrost Plus Adhesion Microscope slides and washed three times for 5 min in 1x PBS at room temperature to remove OCT.

AFM measurements. The AFM-based nanoindentation measurements were carried out using an commercial AFM (JPK Nanowizard) equipped with a Nikon optical microscope. AFM nanoindentation tests were performed using a 5-μm-radius cylindrical tipped nitride cantilever (SAA-SPH-5UM, Bruker) in 1x PBS. Cantilever spring constants were calibrated each time before sample measurements using the thermal fluctuation method, which were in the range of 0.20−0.30 N $m^{-1}$. During measurements, samples were maintained in 1x PBS. Brightfield and K14rt-H2B-GFP images were captured, and used to align the cantilever to the sample and for image co-registration. Two-dimensional force maps were taken in 15 μm × 15 μm square grids with 64 sample points per axial dimension. AFM measurements were made using a cantilever deflection set point of 2.0 nN and an indentation rate of 5 μm $s^{-1}$ to capture elastic properties and minimize viscoelastic effects.

The force-indentation traces were analyzed to obtain the Young's modulus of the cells by using the JPK Data processing program. After baseline correction and contact point estimation, the approaching force-indentation curve was fitted with the Hertz (Spherical) model (eq. 1). Constant parameters were chosen to minimize the bias for different samples.

$$F(x) = \frac{4}{3}\frac{E}{(1-v^2)}\sqrt{r}x^{3/2} \qquad (1)$$

where $F$ is the force of the cantilever, $x$ is the indentation distance of the cell pressed by the cantilever, $E$ is the Young's modulus of the cell layer, $r$ is the radius of the spherical indenter, and

$\upsilon$ is the Poisson ratio. The Poisson ratio of cell is normally in the range of 0.3-0.5. We chose $\upsilon = 0.5$ in all calculations.

**Quantification and statistical analysis**

Quantification and 3D F-actin vs YAP analysis

To quantify the immune-staining signals of YAP and cortical F-actin in hair germs, the images were converted to Imaris for further quantification. Briefly, Imaris Surface module was applied to automatically select YAP positive areas in the nucleus in hair germ cells. The mean/median intensity of YAP signals of the selected region was obtained from the statistical tab. Cortical F-actin signals were quantified using Line quantification, crossing F-actin bundles, in Fiji software. The mean peak intensities of 5 line quantifications per cell was calculated.

After quantification of F-actin and YAP signals in wildtype control and miR-205 ind samples, we did normalization of F-actin values and YAP values by dividing all the values with the maximum value of the condition. For wildtype data, we also assigned a phase of the hair cycle (Telogen and Anagen) to each cell depending on their morphology and hair cycle stages. We then defined Telogen Center, *t*, and Anagen Center, *a*, by calculating the mean of normalized F-actin value and YAP value for Telogen and Anagen cells in wildtype respectively. We then defined a binary phase score for a cell *c* as:

$$\text{Binary Phase Score} = \mathrm{dist}(c, a) - \mathrm{dist}(c, t)$$

, where the distance is between normalized F-actin and YAP of the cell and two centers. Observe that if a cell is closer to the Telogen Center, it will have a positive binary phase score. Otherwise, if it is closer to the Anagen Center, it will have a negative binary phase score. If a cell is equidistant from both centers, it will have a zero score.

For visualization, we defined a Telogen-Anagen phase landscape by finding binary phase scores for simulated normalized F-actin and YAP value combinations ($n=21^2$, where both values in any combination is out of a sequence ranging from 0 to 1, incremented by 0.05) and created a smoothed 3D surface from simulated combinations with

$$\mathrm{x} = \text{normalized YAP value}$$

$$y = \text{normalized F-actin value}$$
$$z = \text{binary phase score}$$

Later, for each cell in control and miR-205 ind samples, we again calculated its binary phase score based on the normalized YAP and F-actin values and then plotted the data to the Telogen-Anagen phase landscape. For each group of data, one-sided Wilcoxon test was used to test the difference in mean.

To investigate the significance of the correlation between YAP and cortical F-actin (PHALL signal), we conducted a randomization test by shuffling all YAP and PHALL value pairs 1000 times, and we calculated R^2 with 1000 for each shuffle and built a background distribution from these 1000 R^2. Finally, we defined the empirical p-value of R-squared as:

$\mathrm{p-value}_{emperical} = \frac{\text{\#of shuffles with } R^2 > R^2_{\mathrm{YAP-PHALL}} + 1}{1000+1}$, where 1 is a pseudocount.

For other cortical F-actin signal quantification, over 10 line quantifications were used for the one hair germ area and they were aligned by the peak values of those line quantification. The mean intensity of F-actin signals for each pixel were used for quantification.

#### Figure S1

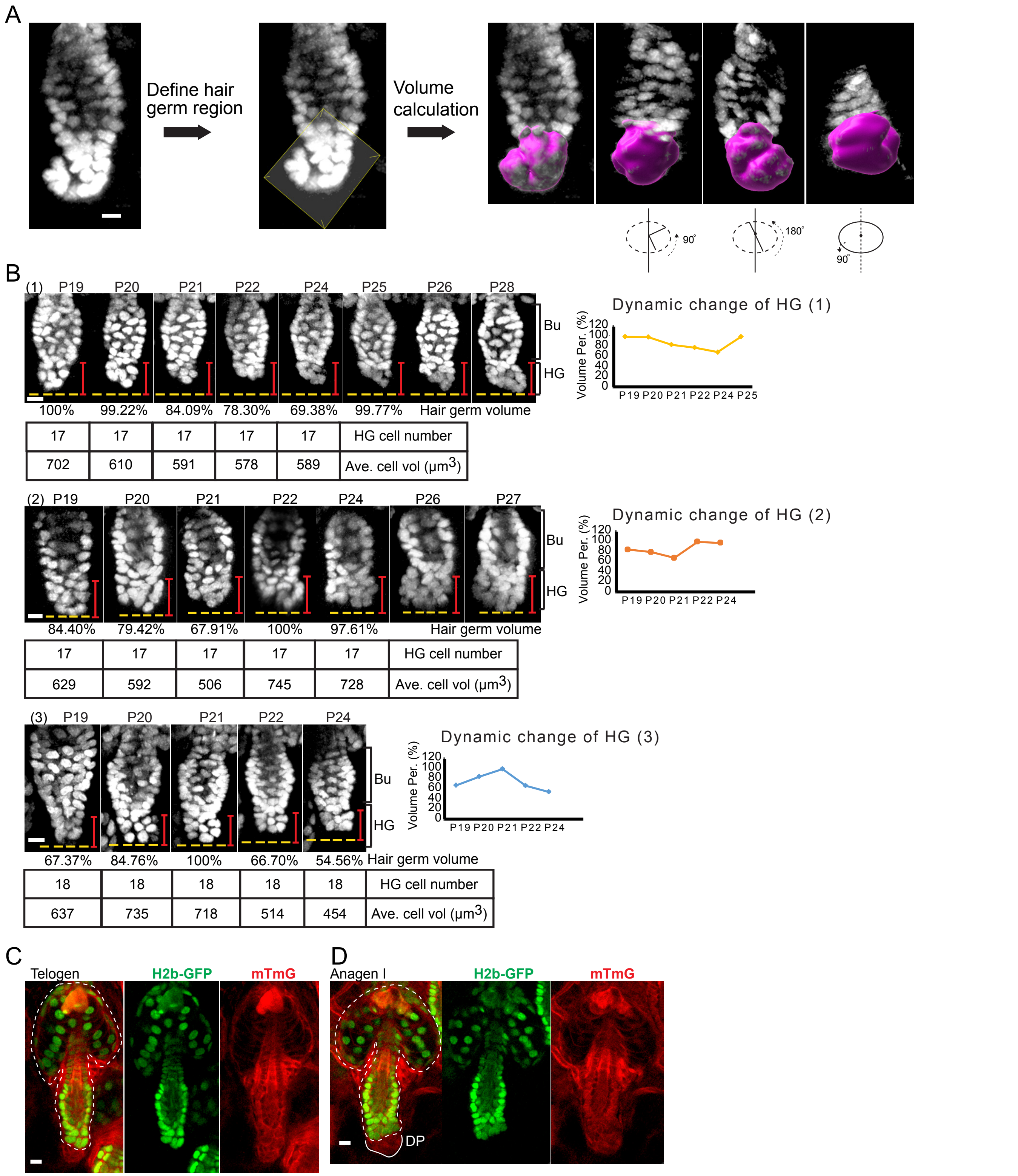


**Figure S1. Quantification of hair germ volumes in live animals.**
(**A**) The workflow for the quantification of HG volume by using H2b-GFP in Imaris. (**B**) Hair germs show dynamic size changes in late catagen and telogen without cell division. The average cell size was calculated for hair germ cells during telogen. Bu, bulge; HG, hair germ. 65 hair follicles from 4 animals were tracked. (**C**-**D**), Double labeling of HF-SCs and HG progenitors by H2b-GFP and mT in telogen and early anagen. DP, dermal papillae.
Scale bar 10 μm in **A**, **C**, **D**, 15 μm in **B**.

#### Figure S2

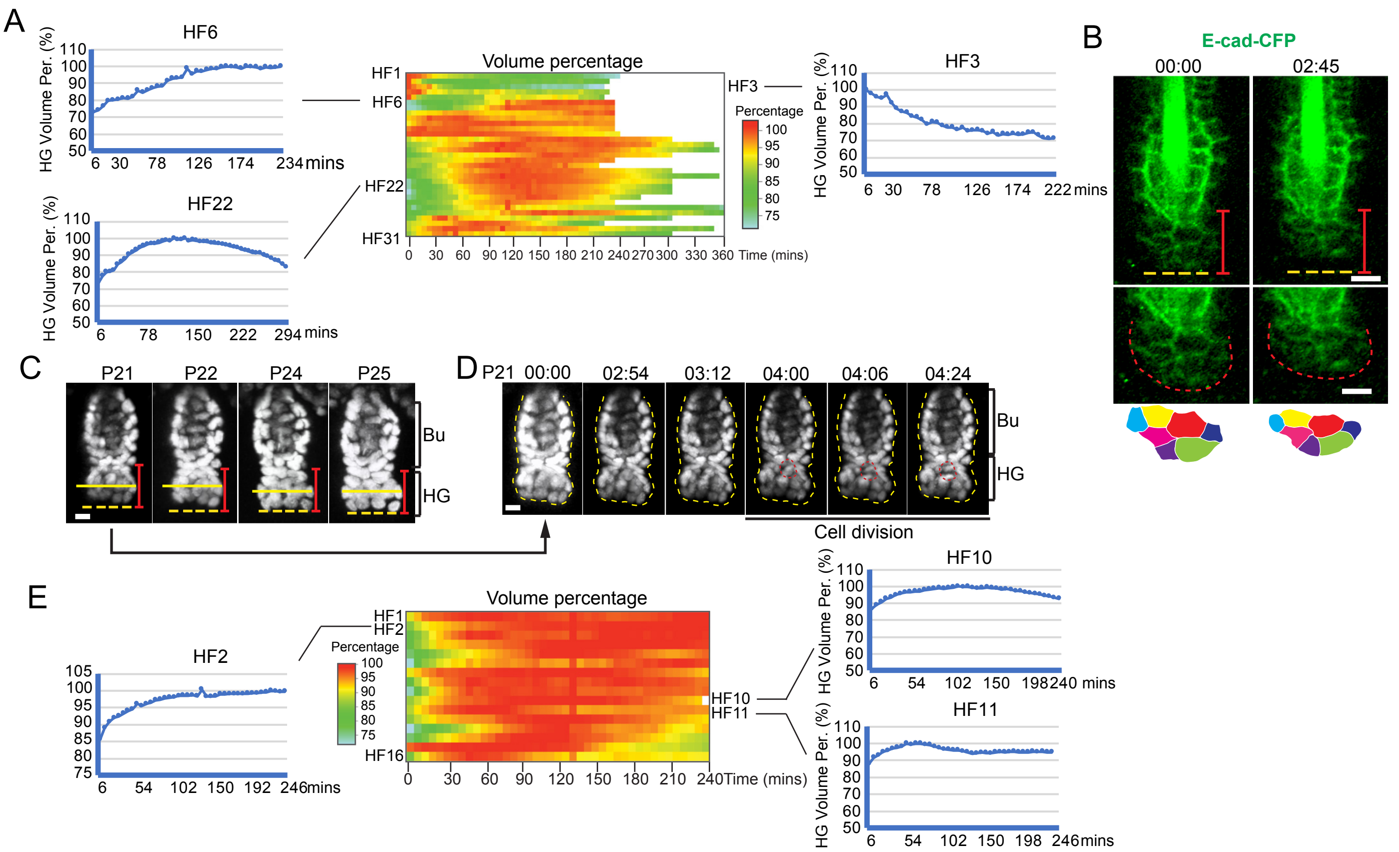


**Figure S2. Hair germs are mechanically active and exhibit contraction and enlargement.** (**A**) HGs in telogen exhibit cyclic contraction and enlargement. 3 different patterns were shown as examples. 31 hair follicles from 9 animals were used to quantify volume changes. (**B**) HG contraction during telogen was recorded by visualizing E-Cad-CFP signals at the cell membrane. (**C**) An example of HG undergoing enlargement during early anagen. (**D**) Infrequent cell division was recorded in an HG at early anagen(P21). No HG volume changes were recorded during the division. (**E**) HGs in early anagen exhibit reduced contraction, comparing with those in telogen. 16 hair follicles from 5 animals were used to quantify volume changes.
Scale bar 10 μm in **B**, **C**, **D**.

**Figure S3**

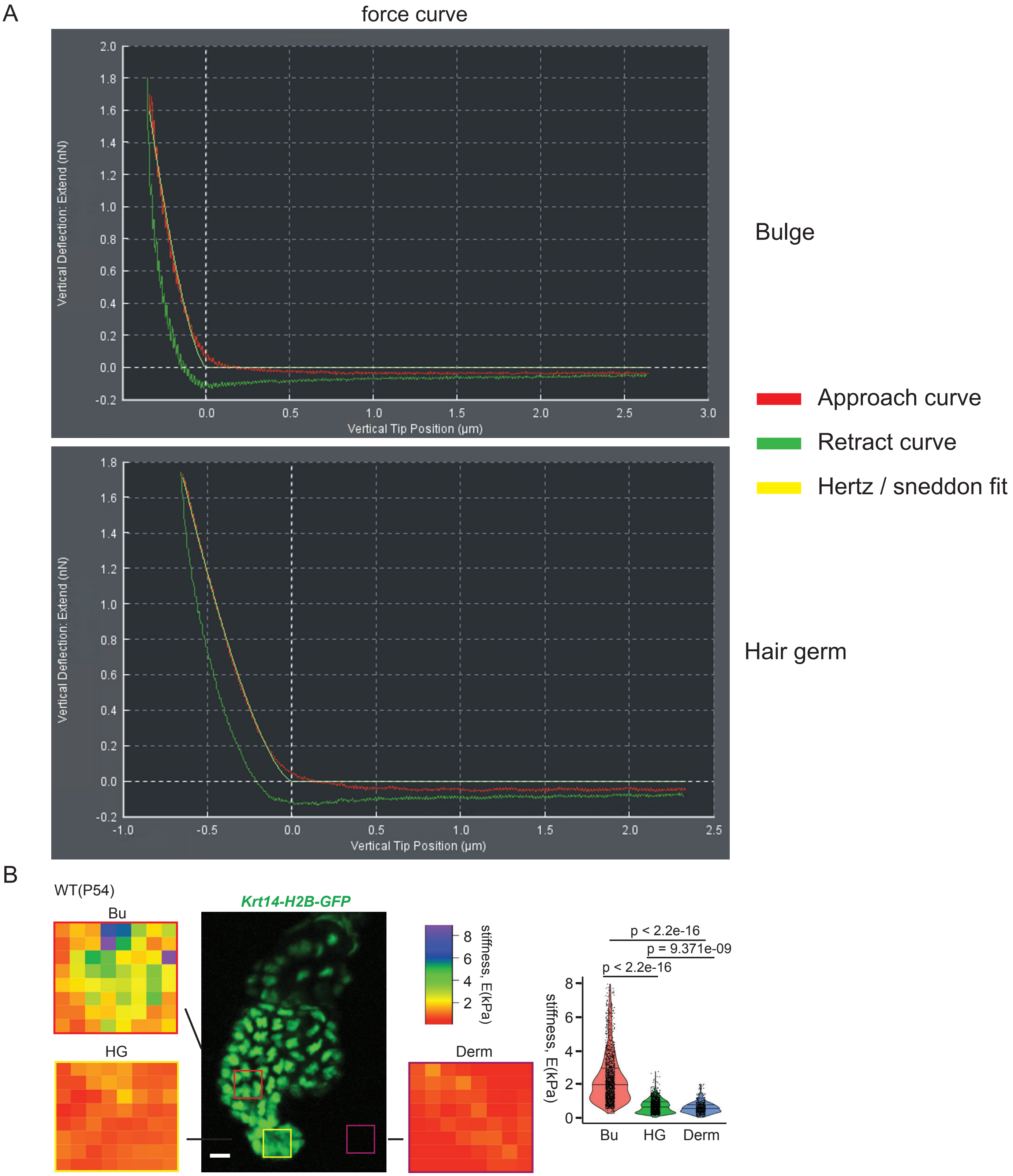


**Figure S3. HF-SC compartment is demarcated by differential mechanical stiffness measured by AFM**
(**A**) Force curve and Hertz / Sneddon fit for Young's modulus measurement of the bulge and HG.
(**B**) Stiffness of the bulge, HG and dermis was measured ex vivo by atomic force microscopy (AFM). Six hair follicles and dermis areas from 2 animal samples were measured. For each area, 64 points were measured.
Bu mean= 2240.1 ± 1251.1 Pa, HG mean= 824.2 ± 368.0 Pa, Dermis mean= 617.3 ± 329.3 Pa.
P values were determined by Student's t-test. Scale bar, 10 µm.

#### Figure S4

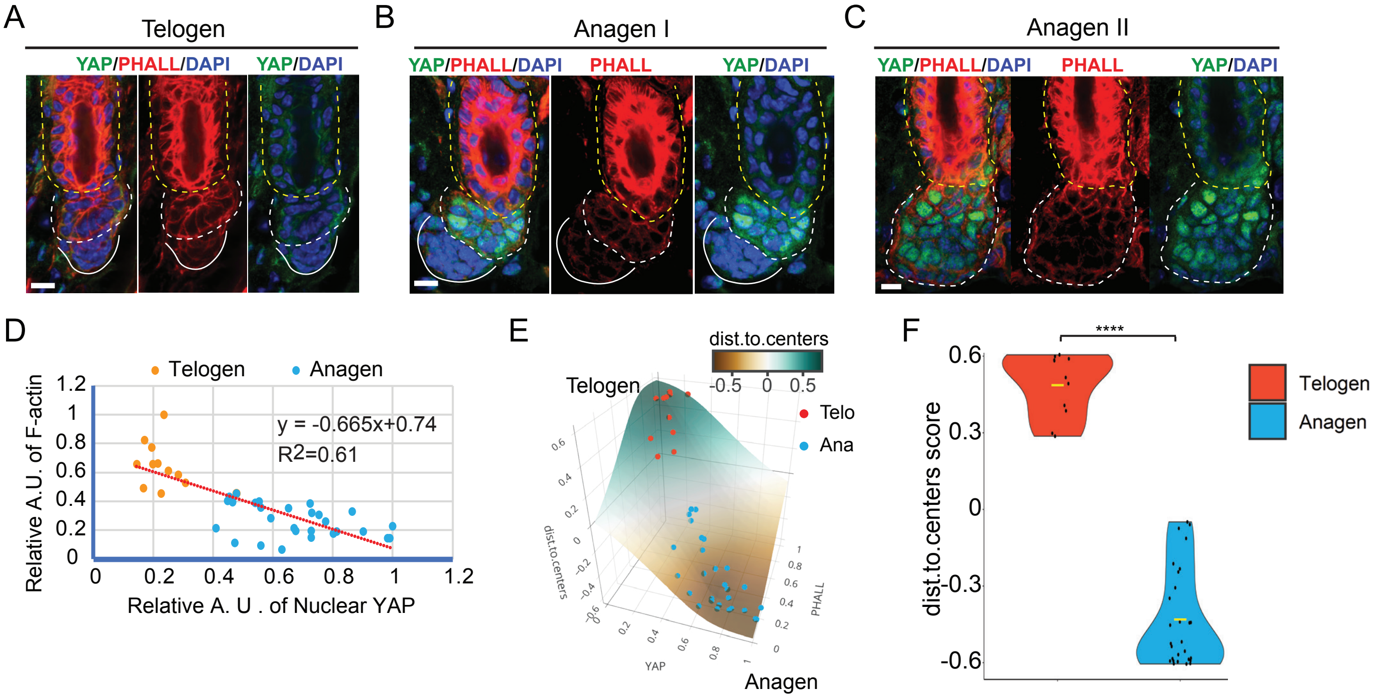


**Figure S4. Weakened actomyosin contractility is correlated with nuclear YAP accumulation during hair germ progenitor activation**

(**A-C**) Representative confocal microscopy image of telogen (**A**), anagen **I** (**B**) and anagen **II** (**C**) hair follicles with YAP and F-actin staining. (**D**) Cortical F-actin and nuclear accumulation of YAP showed an inverse correlation in telogen and early anagen. (**E**) Combined F-actin and nuclear YAP score demarcates telogen and anagen cellular states, as plotted in a 3-dimensional space. The color indicated the “distance to centers” score of each sample point. (**F**) F-actin and nuclear YAP signals, modeled by a distance to centers score, demarcate the telogen and anagen state (****p<0.0001).

Scale bar 10 μm in **A**, **B**, **C**.

### Figure S5

A

**miR-205 bound components of actin cytoskeleton and cell adhesion captured by CLEAR-Clip**

| **Supramolecular fiber organization** | |
|---|---|
| Regulation of cytoskeleton organization | Cav1, Cd2ap, Cdkn1b, Il1a, Map1b, Notch2, Pafah1b1, Vps4b, Hdgfl3, Dlc1, Clip1, Cltc, Syde1, Actr3, Slain2, Mtpn, Tmsb4x, Hdgfl3, Kank1, Arfgef1, Taok1, Ppfia1 |
| Actomyosin structure organization | Cflar, Ccn2, Mef2a, Met, Dlc1, Tacstd2, Cdc42bpb, Pak2, Ppfia1, Fhod1, Kank2, Epb41, Frmd6 |
| Regulation of actin cytoskeleton | Actb, Actn4, Actn1, Cdc42, Fn1, Itgav, Ppp1r12a, Ppp1cb, Tmsb4x, Sos1, Iqgap1, Arpc1a, Pak2, Pak4, Baiap2 |
| Actin filament organization | Cd2ap, Cdc42, Dlg1, Nedd9, Shroom3, Rufy3, Sh3kbp1, Ttc17, Marcks, Met, Hsp90b1, Pls3, |
| Regulation of actin filament organization | Dlg1, Ccn2, Mtpn, Met, Tmsb4x, Tjp1, Dlc1, Arpc1a, Tacstd2, Actr3, Rictor, Kank1, Kank2, Baiap2, Arfgef1, Pak2, Ppfia1, Fhod1, Rock2 |

| **Regulation of cell adhesion** | |
|---|---|
| Cell-cell adhesion | Ctnna1, Cd2ap, Chl1, Fut9, Lrp6, Ptprs, Stxbp3, Nt5e, Cd164, Cadm1, Perp, Zdhhc21, Mcam, Lpp, Dll1 |
| Regulation of cell adhesion | Adam8, Slc7a1, Bcl6, Cav1, Runx1, Cd44, Cdk6, Fn1, Hes1, Il1a, Il6st, Itgav, Nedd9, Prkca, Cyth3, Ptprj, Sox2, Sox4, Stx3, Stk4, Thbs1, Tjp1, Map4k4, Net1, Piezo1, Zmiz1, Cd9, Dlg1, Jag1, Notch1, Dusp1, Snai2, Tnfaip3, Dlc1, Tacstd2, Peli1, Scrib, Cblb, Acer2, Wnk1 |
| Adherens junction | Ctnna1, Lmo7, Igf1r, Met, Ptprj, Snai2 |
| Focal adhesion | Cav1, Ccnd2, Fn1, Igf1r, Itgav, Met, Ppp1r12a, Ppp1cb, Prkca, Sos1, Thbs1 |
| Tight junction | Runx1, Cldn1, Cldn4, Dlg1, Ppp2cb, Tjp2, Pard6b, Rapgef2, Scrib |

B

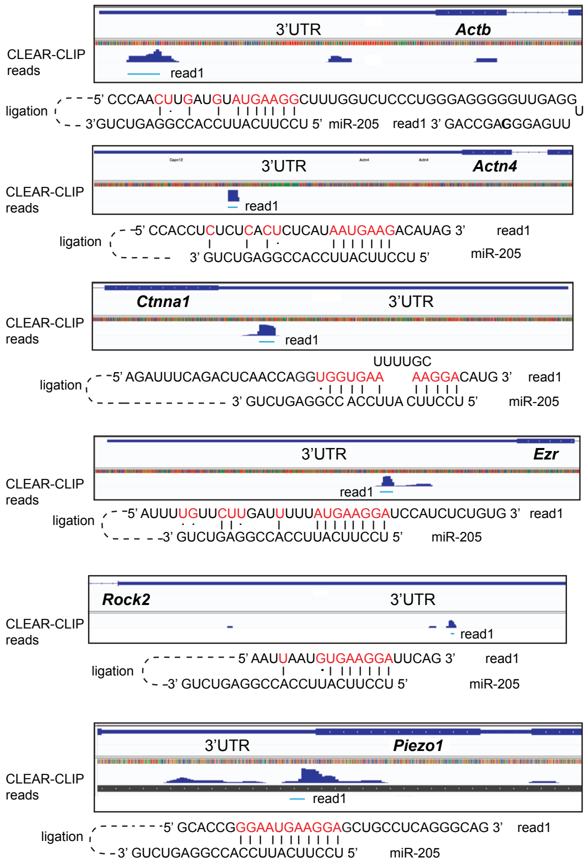


C

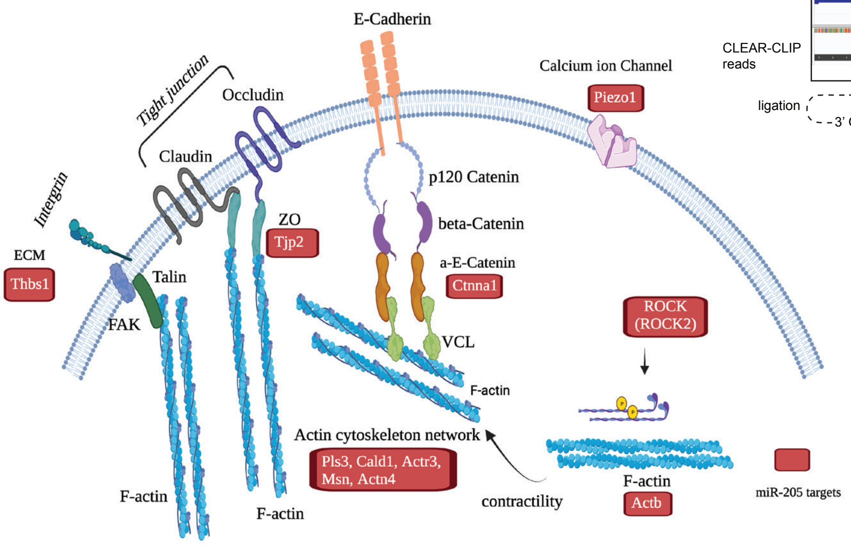


**Figure S5. *miR-205* targets components and regulators of the actin cytoskeleton and cell adhesion machinery**
(**A**) *miR-205* binds to mRNAs of components and regulators of the actin cytoskeleton and cell adhesion, determined by CLEAR-CLIP. Genes marked by red font were highlighted in the manuscript. (**B**) The binding of *miR-205* to target sites, located within the 3'UTR of mRNAs, was caputured by CLEAR-CLIP. Base-pairing between *miR-205* and targeted mRNA sites, marked by red font, is shown. (**C**) Illustration of widespread regulation of the actomyosin network, multiple cell adhesion machinery, including, the adherens junction, tight junction and integrin, as well as Piezo-1 by *miR-205*. Experimentally identified *miR-205* targets are highlighted in red color.

### Figure S6

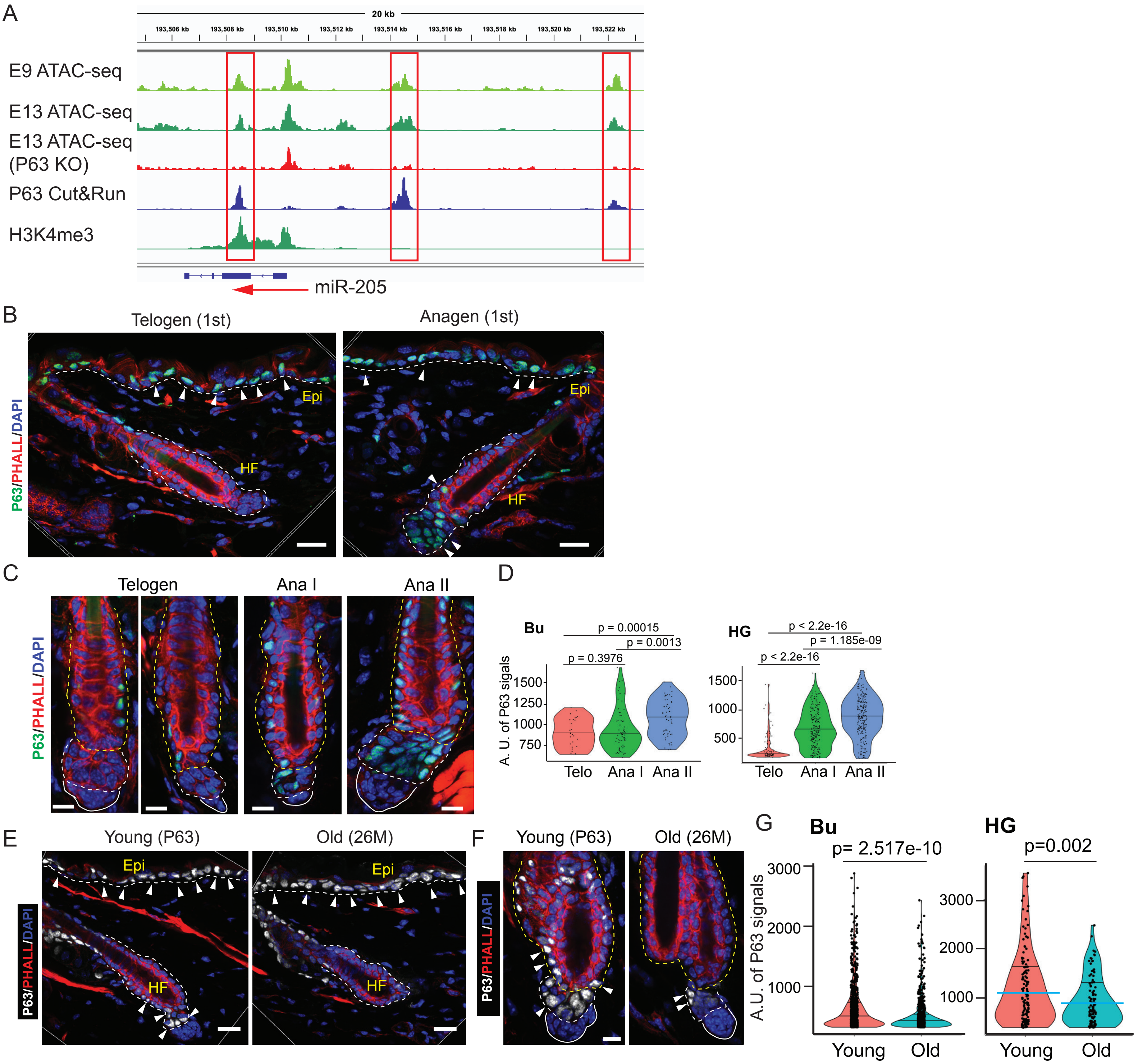


**Figure S6. The expression and regulation of *miR-205* in telogen, early anagen and durinag aging.** (**A**) IGV track of ΔNp63 ATAC-seq datasets shows that *miR-205* is transcriptionally regulated by ΔNp63 through three enhancers, identified by ΔNp63 Cut&Run. Note the open chromatin signals are lost in ΔNp63 KO. (**B**) ΔNp63 shows different expression patterns in bulge and HG in telogen and early Anagen. Note the largley uniform expression of ΔNp63 in the epidermis. Scale bar, 20 μm. (**C**) ΔNp63 expression level is elevated in both bulge and HG in early anagen, compared with that in telogen. Scale bar, 10 μm. (**D**) Quantification of ΔNp63 levels from bulge and HG in telogen and anagen. Nuclear ΔNp63 signals from 6 telogen hair follicles of 3 animals, 7 anagen I hair follicles of 3 animals and 8 anagen II hair follicle from 3 animals are quantified. P values are determined by Student's t-test. (**E**-**F**) Nuclear ΔNp63 expression level is decreased in both bulge and HG in old mice. Note the largley uniform expression of ΔNp63 in the epidermis. Scale bar 20 μm in E, 10 μm in F. (**G**) Quantification of ΔNp63 levels from bulge and HG in young and old dorsal skin. Nuclear ΔNp63 signals from 11 hair follicles of 3 old animals, 16 hair follicles of 3 young animals are quantified. P values were determined by Student's t-test. In **B** and **E,** arrowheads point to ΔNp63 positivie nuclei.

### Figure S7

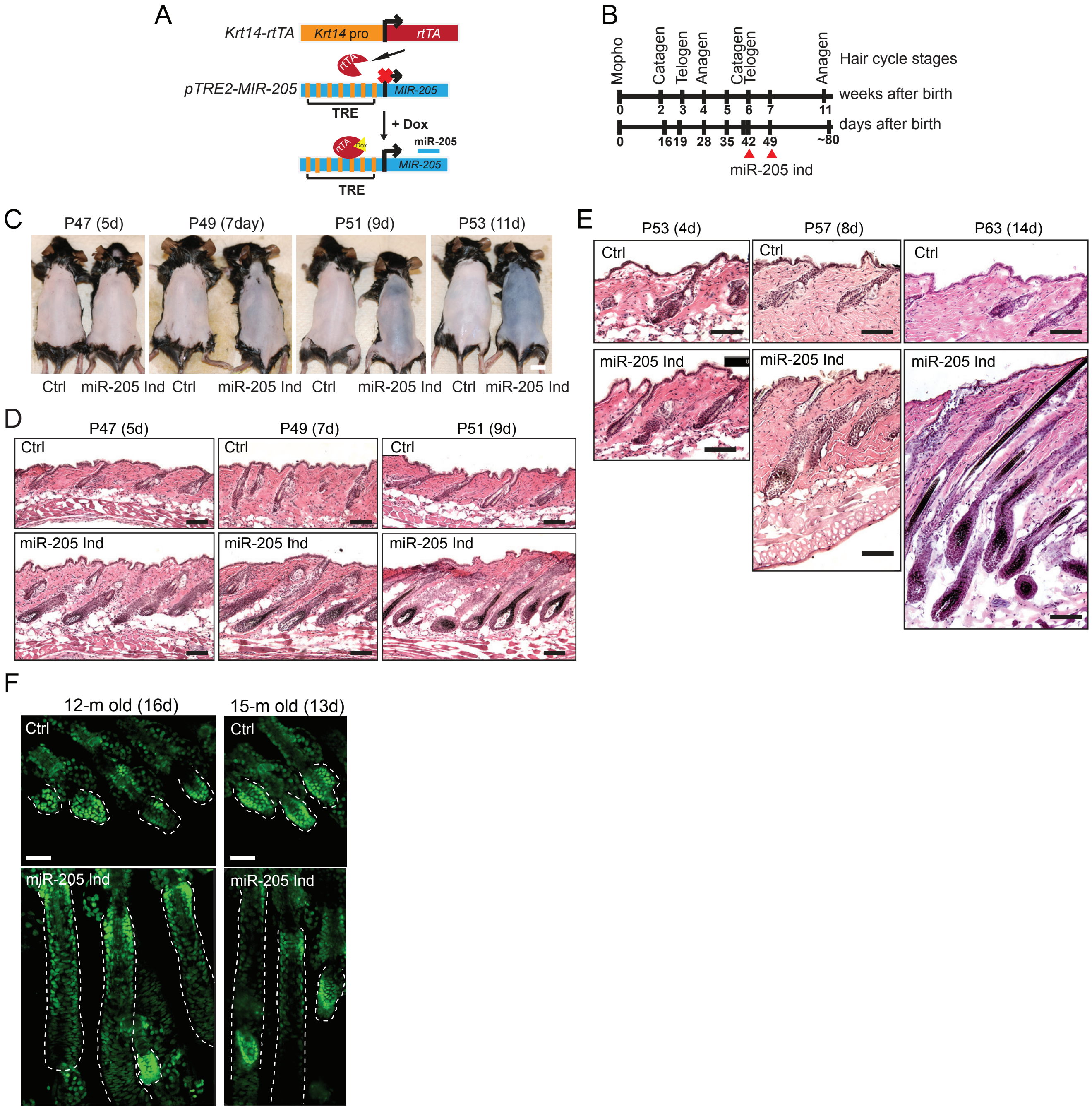


**Figure S7. *miR-205* induction promotes hair regeneration in young and old mice.**
(**A**) Schematics of *miR-205* inducible mouse model driven by *Krt14-rtTA*. (**B**) Schematics of *miR-205* induction in young mice. (**C**) *miR-205* promotes hair regeneration when the induction is initiated at P42. 5 pairs of animals were used for phenotypical analysis at each stage. (**D**) H&E staining for *miR-205* induced skin after 5-d, 7-d and 9-d induction. (**E**) H&E staining for *miR-205* induced skin after 4-day, 8-day and 14-day induction. (**F**) *miR-205* promotes ear hair regeneration in old mice, longitudinally tracked with multiphoton microscopy.
Scale bar 1 cm in **C**, 100 μm in **D**, **E**, **F**.

#### Figure S8

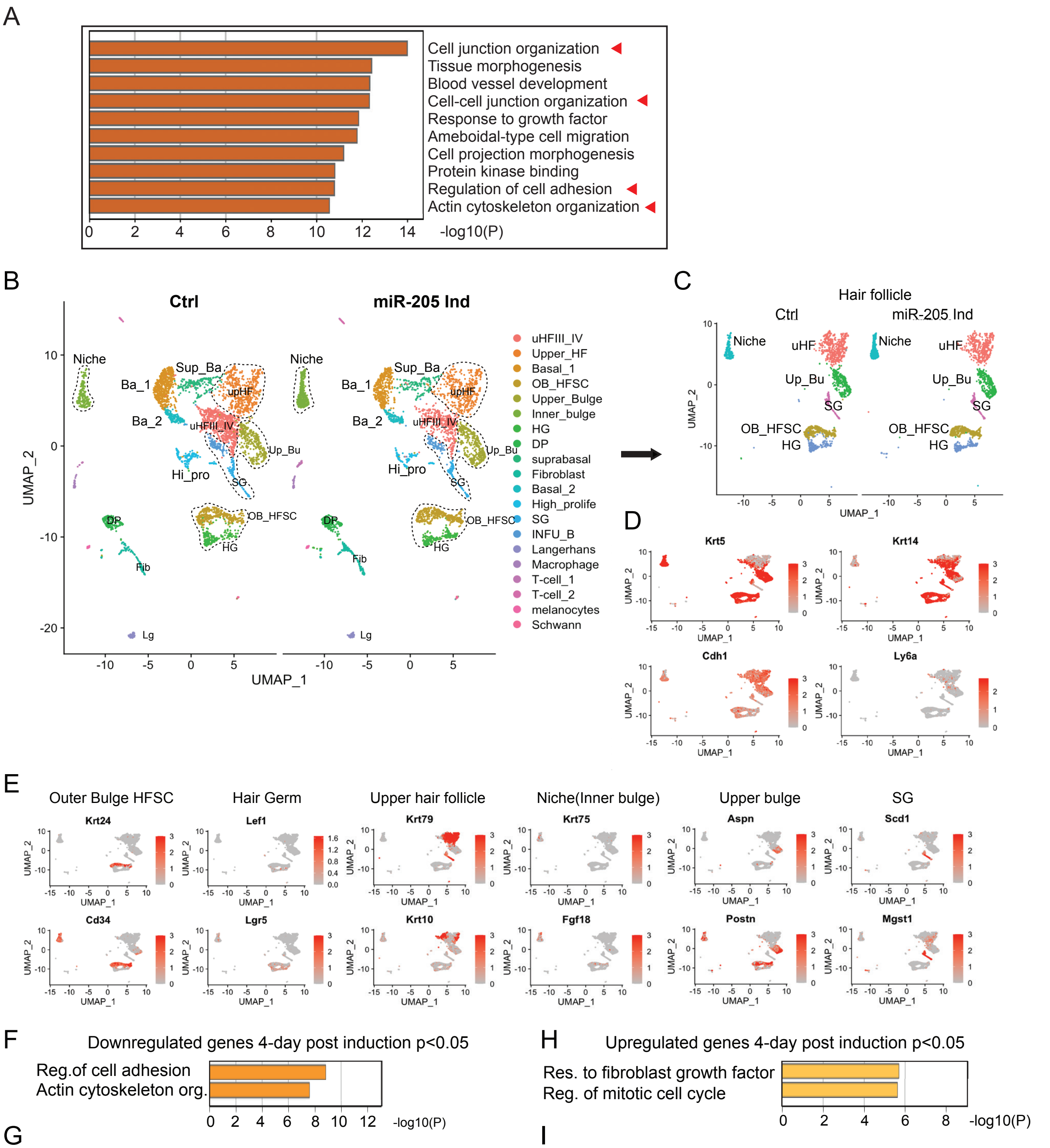


G

| GO Term | Genes (log2FC) |
|---|---|
| **Regulation of cell adhesion** | Cldn1 (-0.27), Tjp2 (-0.72), Epcam (-0.41), Piezo1 (-0.72), Nedd9 (-0.47), Cadm1 (-1.12), Itgb1 (-0.68), Cd44 (-0.46), Thbs1 (-1.36), Crb3 (-0.86), Itga2 (-0.65), Itga3 (-0.56), Cd63 (-0.44), Epha2 (-0.44), Adgrg1 (-0.73), Has2 (-0.78), Tnc (-0.55), Vcl (-0.46), Lgals3 (-0.87), Lgals7 (-0.30), Ccn1, Zyx (-0.37), |
| **Actin cytoskeleton organization** | Actb (-0.54), Cfl1 (-0.44), Anxa1 (-0.49), Vasp (-0.49), Tacstd2 (-0.63), Macf1 (-0.47), Actg1 (-0.57), Capg (-0.57), Capza2 (-0.33), Enah (-0.44), Marcksl1 (-1.10), Pak2 (-0.71) Rock1 (-0.44), Pdlim1 (-0.35), Palld (-0.74)<br>Actin-membrane linkers :Msn (-0.50), EZR (-0.32), Iqgap1 (-0.32), UTRN (-0.51)<br>Actin crosslinker: Actn1 (-0.54), Actn4 (-0.49), Pls3 (-0.49), Flnb (-0.62), TPM3 (-0.33), TPM1 (-3.88)<br>Actin filament nucleators:Actr3 (-0.43), Arpc2 (-0.49), Arpc3 (-0.33)<br>Contractility: Pak1(-0.63), Rock2 (-0.45), MYL12A |

I

| GO Term | Genes (log2FC) |
|---|---|
| response to fibroblast growth factor | Fgfr1 (0.52), Fgfr3 (1.33), Tbx1 (0.82) |
| regulation of mitotic cell cycle | Cdc25b (1.39), Mki67 (3.62), Tcf3 (0.68), Wee1 (0.88), Cdc23 (1.49), Mad2l1bp (2.62), Nabp2 (1.31), Bora (2.20), Scrib (1.28), Cenpf (3.90), Pim3 (0.70), Cenpe (5.16), Npm1 (0.55), Psrc1 (4.80), Ccni (0.63), Eps8 (0.80), Tcim (1.99) |
| Wnt signaling pathway | Dlx5 (1.85), Egr1 (1.08), Fzd1 (1.58), Wnt4 (1.28), Sostdc1 (0.95), Wls (0.55), Trpm4 (1.97), Fam53b (1.59) |

**Figure S8. *miR-205* targets components and regulators of the actin cytoskeleton and cell adhesion machinery in HF-SCs**

(**A**) *miR-205* targets in the gene categories of the actin cytoskeleton and cell junction are the most highly enriched gene groups and are downregulated in HF-SC cells after 2-day induction. (**B**) *miR-205* induction does not globally change the epithelial and dermal cell fate 4-day after induction. (**C**) *miR-205* does not globally change the epithelial cell fate 4-day after induction. (**D**) Epithelial cells are identified by established marker genes *Krt5*, *Krt14* and *Cdh1*. (**E**) Epithelial cell clusters are identified by established marker genes. (**F**) Genes associated with the regulation of cell adhesion and actin cytoskeleton organization are the most highly downregulated genes in *miR-205* induced HGs 4 days after induction. (**G**) Downregulated genes in the categories of of cell adhesion and actin cytoskeleton organization in *miR-205* induced HGs 4 days after induction. *miR-205* targeted genes are highlighted with red color. (**H**) Genes associated with mitotic cell cycle, FGF and WNT signaling are among the most highly upregulated genes in *miR-205* induced HGs 4 days after induction. (**I**) Upregulated genes in the categories of mitotic cell cycle, FGF and WNT signaling in *miR-205* induced HGs 4 days after induction.

**Figure S9**

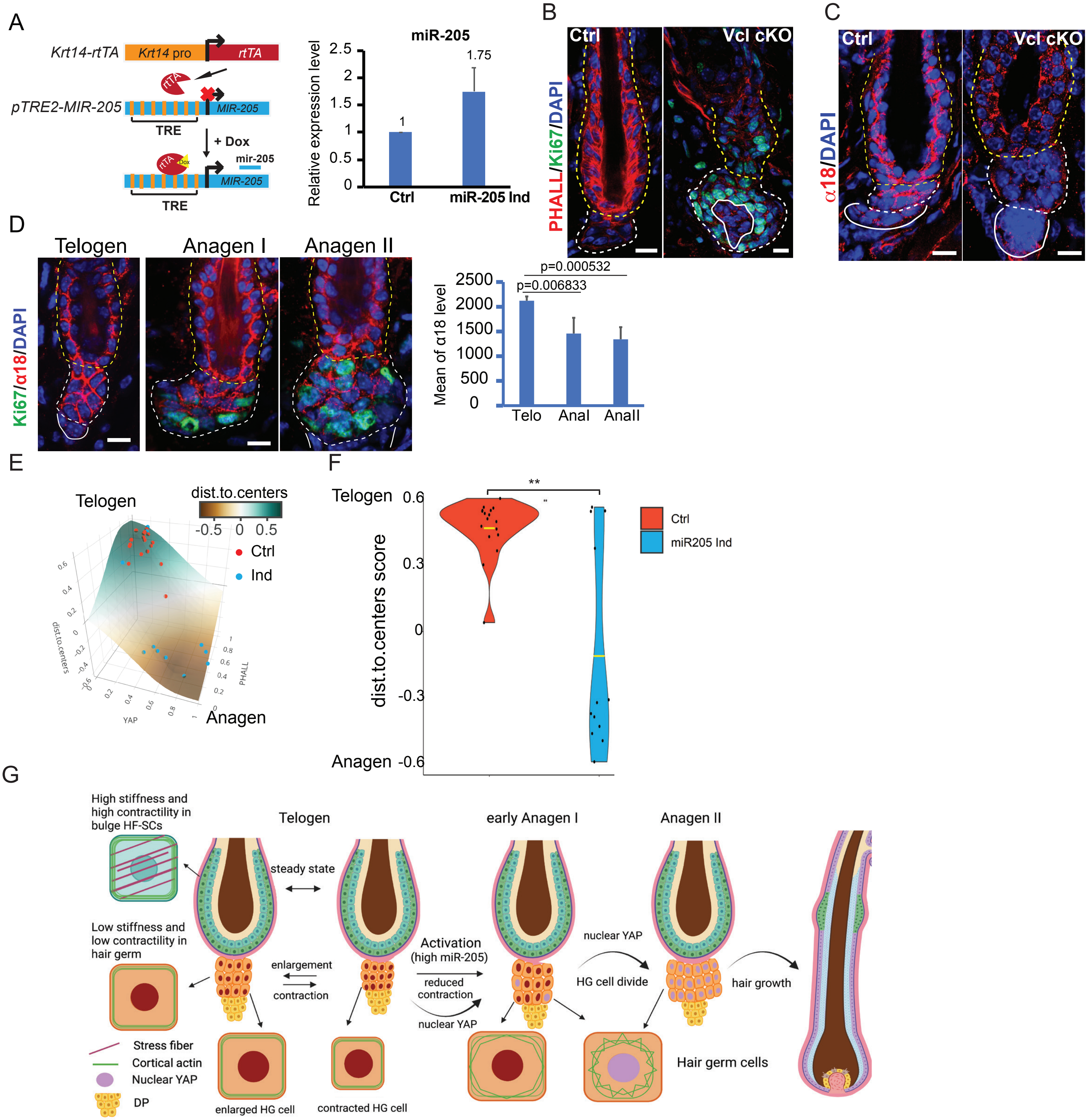


**Figure S9. Interplay between tissue stiffness, actomyosin contractility and cell size control during the transition between quiescence and activation of hair regeneration.**

(**A**) *miR-205* levels are elevated by 75% upon 24-h induction in primary keratinocytes. (**B**) *Vcl* deletion reduces actin cytoskeleton and promotes cell cycle re-entry in bulge HF-SCs and HGs (n=3 pairs of animals). (**C**) Actomyosin forces on α-catenin are significantly reduced upon Vcl loss, indicated by reduced α18 signals (n=3 pairs of animals). (**D**) Actomyosin forces on α-catenin are stronger in telogen than in early anagen, indicated by reduced α18 signals. Quantification is showed in the right panel. P values are determined by Student's t-test. (**E**) *miR-205* induction promotes the transition of HG cells from the quiescent telogen state to the activated anagen state, visualized by mathematical modeling and plotting of the F-actin and nuclear YAP score. The color gradient indicates the “distance to centers” score of each sample point. (**F**) F-actin and nuclear YAP signals, modeled as a distance to centers score, indicate that *miR-205* induction promotes the transition of HG cells from telogen to anagen. Note some cells are still classified as telogen cells whereas many are classified as anagen (**p<0.01). (**J**) Schematic illustration of the interplay between tissue stiffness, actomyosin contractility, nuclear YAP accumulation and cell size dynamics during the transition between quiescence and activation of hair regeneration.

**Dataset S1.**

Differential Gene Expression Upon miR205 Induction (2 days) in HFSCs, determine by bulk RNA-seq

**Dataset S2.**

Differential Gene Expression Upon miR205 Induction (4 days) in HFSCs and HG, determined by single-cell RNA-seq

**Movie S1. Hair germ contraction during telogen.**

In the 4-hour intravital imaging, the HG but not the bulge HF-SC compartment contracts by ~30% in volume. Note the constant position and location of each HF-SCs during the same period. All epithelial cells are labeled with H2b-GFP. The constant red dotted line outlines the HF-SC and HG compartments. The yellow dotted line and green arrowheads track the movement of the bottom of the HG.

**Movie S2. Dynamic contraction and enlargement activities of hair germ during telogen.**

In the 6-hour intravital imaging, the HG exhibits pulsatile contraction and enlargement activities. Note the constant position and location of HF-SCs during the same period. All epithelial cells are labeled with H2b-GFP. The green arrowheads track the movement of the bottom of the HG.

**Movie S3. Hair germ contraction during telogen.**

In the 4-hour intravital imaging, the HG but not the bulge HF-SC compartment contracts by ~30% in volume. The plasma membrane of epithelial cells is labeled with E-Cad-CFP. Note the strong E-Cad levels in the HF-SCs and the relatively weak E-Cad levels in the HG. The constant red dotted line outlines the HF-SC and HG compartments. The yellow dotted line and green arrowheads track the movement of the bottom of the HG.

**Movie S4. Hair germ enlargement during early anagen.**

In the 4-hour intravital imaging, the HG enlarges by ~20% in volume in the absence of cell division. Note the HF-SC compartment also enlarges but less than the HG. All epithelial cells are labeled with H2b-GFP. The constant red dotted line outlines the HF-SC and HG compartments. The yellow dotted line and green arrowhead track the movement of the bottom of the HG.

**Movie S5. Hair germ enlargement in an miR-205 induced hair follicle.**

In the 4-hour intravital imaging, the HG initially enlarges and then slightly contracts toward the end of the imaging session in the absence of cell division. Note the HF-SC compartment also enlarges slightly. All epithelial cells are labeled with H2b-GFP. The constant red dotted line outlines the HF-SC and HG compartments. The yellow dotted line and green arrowheads track the movement of the bottom of the HG.
